# Multimodal framework to resolve variants of uncertain significance in *TSC2*

**DOI:** 10.1101/2024.06.07.597916

**Authors:** Carina G Biar, Cole Pfeifer, Gemma L Carvill, Jeffrey D Calhoun

## Abstract

Efforts to resolve the functional impact of variants of uncertain significance (VUS) have lagged behind the identification of new VUS; as such, there is a critical need for scalable VUS resolution technologies. Computational variant effect predictors (VEPs), once trained, can predict pathogenicity for all missense variants in a gene, set of genes, or the exome. Existing tools have employed information on known pathogenic and benign variants throughout the genome to predict pathogenicity of VUS. We hypothesize that taking a gene-specific approach will improve pathogenicity prediction over globally-trained VEPs. We tested this hypothesis using the gene *TSC2*, whose loss of function results in tuberous sclerosis, a multisystem mTORopathy affecting about 1 in 6,000 individuals born in the United States. *TSC2* has been identified as a high-priority target for VUS resolution, with (1) well-characterized molecular and patient phenotypes associated with loss-of-function variants, and (2) more than 2,700 VUS already documented in ClinVar. We developed <u>T</u>uberous sclerosis classifier to <u>R</u>esolve variants of <u>U</u>ncertain <u>S</u>ignificance in *<u>T</u>SC2* (TRUST), a machine learning model to predict pathogenicity of *TSC2* missense VUS. To test whether these predictions are accurate, we further introduce curated loci prime editing (cliPE) as an accessible strategy for performing scalable multiplexed assays of variant effect (MAVEs). Using cliPE, we tested the effects of more than 200 *TSC2* variants, including 106 VUS. It is highly likely this functional data alone would be sufficient to reclassify 92 VUS with most being reclassified as likely benign. We found that TRUST’s classifications were correlated with the functional data, providing additional validation for the *in silico* predictions. We provide our pathogenicity predictions and MAVE data to aid with VUS resolution. In the near future, we plan to host these data on a public website and deposit into relevant databases such as MAVEdb as a community resource. Ultimately, this study provides a framework to complete variant effect maps of *TSC1* and *TSC2* and adapt this approach to other mTORopathy genes.

## Introduction

Clinical genetic sequencing is now routine for epilepsy of suspected genetic etiology. One of the key signaling pathways implicated in epilepsy is the mammalian target of rapamycin (mTOR) signaling pathway. Pathogenic variants in at least 16 mTOR pathway genes result in mTORopathies, a group of neurodevelopmental disorders commonly associated with epilepsy, in both children and adults.^1^ Variants in mTOR pathway genes such as *MTOR* or *DEPDC5* also cause more than 10% of all focal epilepsies.^2^ The mTORopathies are collectively defined by constitutive activity of mTOR complex 1 (mTORC1). In the most common mechanism of pathogenesis, loss-of-function (LOF) variants in mTOR inhibitors (e.g., *TSC1, TSC2, SZT2*, and *DEPDC5*) lead to disinhibition of MTOR and, accordingly, constitutive mTORC1 activation. Causal variants have been associated with both autosomal dominant and autosomal recessive modes of inheritance, and both mendelian patterns exhibit incomplete penetrance.^3^ mTORopathies are also associated with a vast array of comorbidities and presentations, with common features including cytomegaly, neuronal hyperexcitability, and structural brain abnormalities.^4^ Despite these variabilities in genetic mechanism and phenotypic presentation, the convergence on mTORC1 constitutive activity has provided a tractable target for precision medicine development, and several mTOR inhibitors have been trialed in multiple conditions.^5^

The number of variants of uncertain significance (VUS) reported per year is consistently increasing for epilepsy-related genes, including mTORopathy genes. In 2019, a cohort of 9,769 individuals were sent for epilepsy panel testing, resulting in >16,000 reported VUS and only 2,101 pathogenic or likely pathogenic (PLP) variants.^6^ The same phenomenon is observed for most genes associated with at least one genetic condition; as of early 2024, the number of VUS reported in ClinVar has exceeded 1 million.^7^ This rapid rise in VUS reports has outpaced the scalability of classic, low-throughput functional assays. High-throughput multiplexed assays of variant effect (MAVEs) have emerged as a scalable method with the potential not only to resolve VUS, but also to create an atlas of variant effects for many positions in human and non-human genomes.^8^

mTORopathy genes have a significant burden of VUS that complicate variant interpretation, genetic counseling, and patient management. Herein, we focus on the mTORopathy gene tuberous sclerosis complex 2 (*TSC2*), the highest ranked mTORopathy gene in a study aimed at prioritizing genes for variant effect mapping.^9^ Tuberous sclerosis, a genetic disorder affecting multiple organ systems characterized by epilepsy and benign tumors, is caused by pathogenic variants in two mTOR pathway genes, *TSC1* and *TSC2*.^10^ We employed a multimodal approach incorporating both machine learning and scalable functional characterization with the goal of resolving *TSC2* missense VUS. First, we compared the performance of a gene-specific classifier to the performance of many globally-trained *in silico* pathogenicity prediction algorithms and found that our gene-specific classifier outperforms modern variant effect predictors (VEPs) including AlphaMissense, ClinPred, REVEL, and VARITY.^11-14^ We then adapted our previously published low-throughput VUS resolution assay for the gene *SZT2* to increase scalability and screened more than 200 *TSC2* variants.^15^ Our assay validated well on internal assay controls and a truth set of 25 missense variants of known effect, which enabled us to utilize functional scores at a moderate level of evidence for either benignity or pathogenicity. Overall, we present a multimodal approach to VUS resolution in mTORopathy gene *TSC2* that could serve as a general framework for VUS resolution in other mTORopathy genes.

## Materials and Methods

### Supervised learning

*TSC2* missense variants were curated from a variety of sources including: (1) gnomAD v4, (2) the Regeneron exome server, (3) ClinVar, (4) and the *TSC2* Leiden Open Variation Database (LOVD).^16-19^ We subset this data into a truth set of variants present in ClinVar, containing 246 benign or likely benign (BLB) variants and 130 PLP variants. Variants overlapping this ClinVar truth set were discarded from the gnomAD, Regeneron, and LOVD datasets. Variants identified as recurrent (total allele count greater than 3) in population databases (gnomAD or Regeneron) were considered as likely to be enriched in BLB variants (n=86). Variants manually curated as PLP in the *TSC2* LOVD database were considered as likely to be highly enriched in PLP variants (n=120). This set of unique variants was used as an independent holdout dataset for assessment of machine learning model performance. Each variant was annotated with each feature in the feature set (n=36; Table S1) using Ensemble’s Variant Effect Predictor and custom scripts.^20^ Empty values were replaced with the median value for that feature across the full dataset. Correlations of input features are shown in Figures S1 and S2.

Random forest classifiers were developed and performance examined using the python scikit-learn package.^21^ Briefly, grid search was used in combination with cross-validation (cv=5) to identify models with optimal hyperparameters using a 60%/40% training/test split of the truth set to validate model performance and test for model overfitting to the training split. The best-performing model was then used to score VUS from ClinVar, the holdout dataset described above, as well as all possible *TSC2* coding single nucleotide variants (SNVs). Overall, a total of 11,728 missense variants (10,623 unique amino acid substitutions) were scored (Table S2).

### Cell lines and plasmids

HEK 293T (ATCC® CRL-3216™) cells were maintained in standard growth medium (Gibco DMEM #11995-073) supplemented with 10% FBS (R&D Systems; 50-152-7067) and PenStrep (Gibco #15140122). HAP1s (Horizon #C669) were maintained in standard growth medium (Gibco IMDM #12440061) supplemented with 10% FBS and PenStrep. HAP1 cells were used at low passage (<10) for all experiments to minimize the percentage of cells with diploid genomes.

pCMV-PEmax-P2A-GFP was a gift from David Liu (Addgene plasmid #180020; http://n2t.net/addgene:180020; RRID:Addgene180020).^22^ pEF1a-hMLH1dn was a gift from David Liu (Addgene plasmid #174824; http://n2t.net/addgene:174824; RRID:Addgene174824).^22^ The pU6-tevopreq1-GG-acceptor was a gift from David Liu (Addgene plasmid #174038; http://n2t.net/addgene:174038; RRID:Addgene174038).^23^ BPK1520 was a gift from Keith Joung (Addgene plasmid #65777; http://n2t.net/addgene:65777; RRID:Addgene65777).^24^ Integrity of all plasmids was confirmed by Sanger or long-read sequencing.

### Cloning individual prime editing constructs

We prescreened for active enhanced prime editing guide RNAs (epegRNAs) following Liu lab recommendations.^25^ For the epegRNA screen, a single epegRNA for each exon was cloned into the pU6-tevopreq1-GG-acceptor backbone using standard Golden Gate assembly (Table S9).^26^ For the MAVE, a nicking gRNA-encoding plasmid for each target exon was cloned using an adjusted version of the Zhang lab target sequence cloning protocol.^27^ Briefly, oligonucleotides encoding the nicking gRNA were synthesized (IDT; Table S8), annealed and phosphorylated, and ligated into linearized expression vector BPK1520.

### Cloning epegRNA libraries

Pools of single-stranded DNA oligos encoding epegRNAs were designed and ordered (IDT oPools; Table S6). The oligo pool was made double-stranded and BsaI restriction enzyme recognition sites were appended via PCR (10 cycles). The PCR-amplified oligo pool was then digested with BsaI-HFv2 (NEB #R3733S), according to the manufacturer’s instructions. The destination vector (pU6-tevopreq1-GG-acceptor) was also linearized by restriction digest with BsaI-HFv2 and purified by gel or column purification. The digested oligos and tevopreq vector were ligated using T4 DNA ligase (NEB #M0202S) according to the manufacturer’s instructions. The pool of assembled plasmids was then transformed into OneShot TOP10 chemically competent *E. coli* (Thermo Fisher #C404003). The transformation was plated onto a 10-cm LB agar plate containing ampicillin (100 µg/mL). The bacteria were cultured overnight before the plates of colonies were scraped. The ZymoPURE II plasmid midiprep kit (Fisher #NC0835048) was used to extract plasmid DNA from the pooled colonies. Once library concentrations were determined, plasmid libraries were analyzed via long-read sequencing (Plasmidsaurus) and custom targeted amplicon sequencing for quality control.

### Prime editing in HEK293Ts

The co-transfection of the epegRNA-containing pU6-tevopreq1-GG-acceptor, pCMV-PEmax-P2A-GFP, and pEF1a-hMLH1dn plasmids into HEK293T cells was carried out using the TurboFectin 8.0 transfection reagent (OriGene #TF81005). 100,000 cells were seeded 24 hours before transfection in 24-well plates. Plasmids were co-transfected in the following amounts: 75 ng (pU6-tevopreq1-GG-acceptor), 263 ng (pCMV-PEmax-P2A-GFP), and 132 ng (pEF1a-hMLH1dn). As the PEmax plasmid also encodes GFP, successfully transfected cells were selected using fluorescence-activated cell sorting (FACS). GFP+ cells were re-plated under standard culture conditions for at least 48 hours. Cell pellets were either used immediately for genomic DNA (gDNA) extraction or frozen at –20°C prior to gDNA extraction. gDNA from prime edited cells was screened for editing using PCR amplifying the target exon followed by Sanger sequencing (Table S7).

### Prime editing in haploid HAP1s

The co-transfection of the epegRNA-containing pU6-tevopreq1-GG-acceptor library, pCMV-PEmax-P2A-GFP, pEF1a-hMLH1dn, and nicking gRNA-containing BPK1520 plasmids into HAP1 cells was carried out using the Neon electroporation system (Thermo Fisher #MPK5000). For co-transfection, 1 million HAP1 cells were electroporated with the following amounts of plasmid DNA (total of 3000 ng): 451 ng (pU6-tevopreq1-GG-acceptor), 1579 ng (pCMV-PEmax-P2A-GFP), 789 ng (pEF1a-hMLH1dn), and 180 ng (BPK1520). Successfully transfected cells were selected using FACS sorting for GFP+ cells. GFP+ cells (typically 15-30%) were re-plated under standard culture conditions until at least 1 million cells were obtained.

### P-S6 FACS

HAP1 cells were washed twice with DPBS and serum-starved overnight by incubation in low-sera media (IMDM + 0.2% FBS). Serum starvation has been previously utilized in low-throughput experiments to establish that loss of *TSC2* function results in constitutive mTORC1 activity.^28, 29^ Starved HAP1 cells were washed twice with DPBS supplemented with 1% v/v Phosphatase Inhibitor Cocktail 3 (Sigma #P0044) before trypsinization with trypLE (trypLE; 5 min; Gibco #25300062) supplemented with 1% v/v Phosphatase Inhibitor Cocktail 3. Cells were then centrifuged and resuspended in BD Fixation/Permeabilization solution (BD Biosciences #554714). After incubating in the fixation solution on ice for 20 minutes, the HAP1 cells were washed twice with BD permeabilization/wash buffer. The CST Alexa594 conjugated rabbit anti-phospho-S6 antibody (#5018; D68F8) was then used to immunolabel P-S6. Cells were incubated with the antibody (1:50 dilution) for 30 minutes on ice in the dark. This antibody had been previously validated for the functional characterization of variants in mTORopathy gene *SZT2*.^15^ Immunolabeled cells were then washed twice more with BD permeabilization/wash buffer before they were flow sorted by P-S6 level on the BD FACSMelody 3-laser cell sorter. A minimum of 200,000 cells in the upper quartile for P-S6 were collected. Unsorted cells were also saved for downstream processing.

### Amplicon sequencing and data analysis

The gDNA of the fixed, FACS-sorted HAP1 cells was extracted using the PureLink™ Genomic DNA Mini Kit (Invitrogen #K182002). Manufacturer’s instructions were followed with the exception of an added incubation (40-60 minutes, 90°C) to reverse formaldehyde-induced crosslinks. The target regions of *TSC2* were then amplified via PCR (Bio-Rad iProof system), followed by an additional barcoding PCR reaction. The locus was then sequenced via short-read amplicon sequencing (Illumina MiniSeq System; 100,000 minimum read-depth). Reads were aligned to the human genome (hg38) and variant allele frequencies were calculated using the jellyfish k-mer counting package.^30^ Variants with allele frequency below 0.1% in unsorted cells were filtered out. Variant allele frequencies were then used to calculate a functional (enrichment) score using the Enrich2 statistical framework.^31^ Functional scores were then normalized across exons as in Buckley et al.^32^ The normalized functional score is referred to as the constitutive mTORC1 activity score (CMAS). All MAVE scores are reported in Tables S4 and S5.

## Results

### TSC2-specific machine learning model outperforms global models on classified variants in TSC2

Data exploration of our *TSC2* variant annotations showed significant differences in feature score distribution between known BLB (class 0) and known PLP (class 1) variants (Figure S3; Tables S1 and S2). For example, MAESTRO, an *in silico* predictor of protein stability, predicts many PLP substitutions impact protein stability while most BLB substitutions do not (Figure S4). We developed <u>T</u>uberous sclerosis classifier to <u>R</u>esolve variants of <u>U</u>ncertain <u>S</u>ignificance in *<u>T</u>SC2* (TRUST), a *TSC2* gene-specific classifier (Random Forest; n_estimators=160; max_depth=3) and then determined the optimal threshold for binary classification of variants in the truth set. TRUST performed well on cross validation (Table S3), which suggested the model was not over-fitting to the training data. TRUST was both specific (91.8% specificity) and accurate (91.4% accuracy) on the test set (Figure 1A); comprehensive model metrics are provided in Table 1. We then investigated the classification performance on an independent holdout dataset. We compared the feature distributions for our holdout dataset relative to the truth set and found that they were quite similar (Figure S5). For example, MAESTRO stability predictions were distributed differently in the holdout set BLB class relative to the PLP class, similar to the truth set (Figure S6). TRUST performed with 89.5% specificity and 83.5% accuracy on the holdout dataset (Figure 1C). We then benchmarked TRUST against a series of globally-trained *in silico* pathogenicity predictors (Figure 1B and 1D; Figures S7 and S8; Tables 1-2). As with TRUST, optimal thresholds for each globally-trained algorithm were determined on the truth set, as pilot evaluation suggested that author-recommended genome-wide thresholds led to poor classification of the *TSC2* truth set. On the test set, the most accurate globally-trained VEPs are ClinPred (91.8% specificity; 92.7% accuracy) and gMVP (93.9% specificity; 93.4% accuracy), which slightly outperform TRUST. On the holdout dataset, gMVP (86.0% specificity and 83.0% accuracy) and AlphaMissense (88.4% specificity and 82.0% accuracy) were both accurate, with model metrics similar to TRUST.

**Table 1:** Machine learning model metrics on test dataset.

| Model | Threshold | Specificity | Recall | Accuracy | Precision | F1 score |
| --- | --- | --- | --- | --- | --- | --- |
| TRUST | 0.4684 | 0.9184 | 0.9057 | 0.9139 | 0.8571 | 0.8807 |
| AlphaMissense | 0.8153 | 0.9286 | 0.8679 | 0.9073 | 0.8679 | 0.8679 |
| BayesDel | 0.3428 | 0.9184 | 0.9057 | 0.9139 | 0.8571 | 0.8807 |
| ClinPred | 0.9456 | 0.9184 | 0.9434 | 0.9272 | 0.8621 | 0.9009 |
| EVE | 0.7068 | 0.8966 | 0.7925 | 0.8571 | 0.8235 | 0.8077 |
| gMVP | 0.749 | 0.9388 | 0.9245 | 0.9338 | 0.8909 | 0.9074 |
| REVEL | 0.788 | 0.8776 | 0.8868 | 0.8808 | 0.7966 | 0.8393 |
| VARITY_R | 0.7563 | 0.9082 | 0.8491 | 0.8874 | 0.8333 | 0.8411 |

**Table 2:**
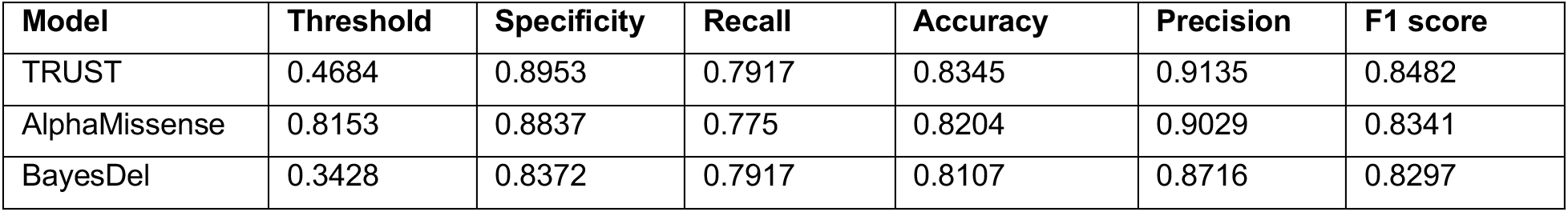
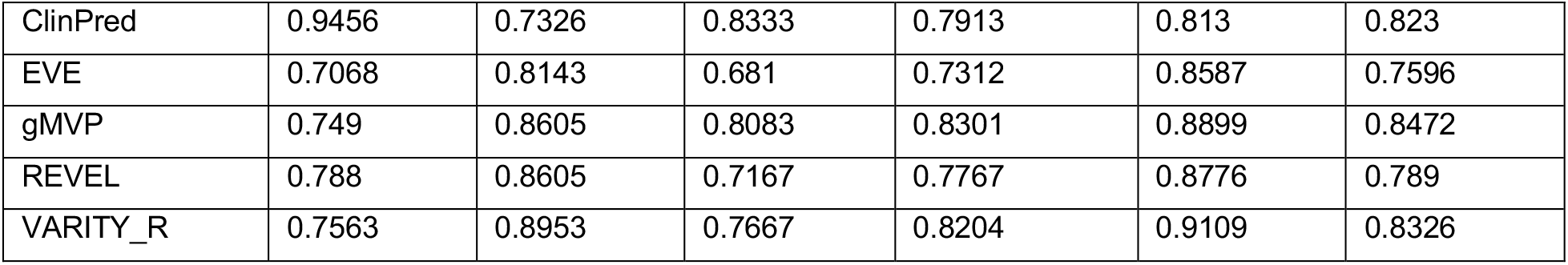
Machine learning model metrics on holdout dataset.

**Figure 1:**
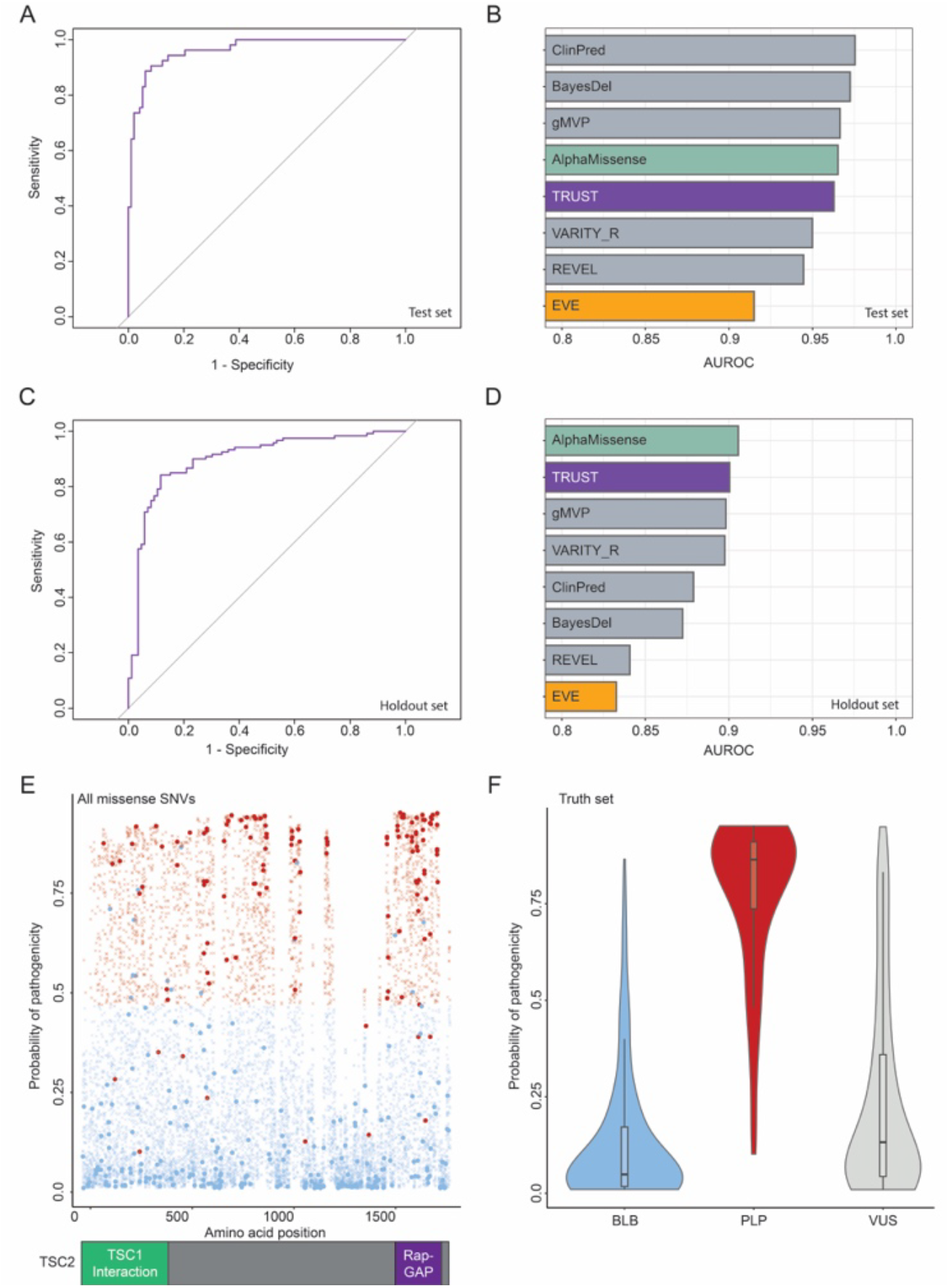
Development and initial evaluation of a *TSC2-*specific classifier for predicting pathogenicity of missense variants. (**A**) Receiver operator characteristic (ROC) curve for TRUST classifier on the test set (n=151 variants). (**B**) Area under the ROC curve (AUROC) for TRUST and some of the best-performing globally-trained VEPs on the test set. Globally-trained VEPs trained on the ClinVar dataset are displayed in gray bars. (**C**) ROC curve for TRUST classifier on the holdout dataset (n=206 variants). (**D**) AUROC for TRUST and some of the best-performing globally-trained VEPs on the holdout dataset. (**E**) Plot of the TRUST output probability of pathogenicity by TSC2 amino acid position for all variants in the study (n=11,728). Truth set variants are displayed as blue (BLB) or red (PLP) circles. All other variants are displayed as either a blue (predicted BLB) or red (predicted PLP) ‘X’ symbol. A cartoon diagram of TSC2 functional domains is provided for reference. (**F**) Violin plot showing the distribution of TRUST probability output by variant class for the full truth set as well as missense VUS.

We next asked if there was a relationship between amino acid position and the classifier output (Figure 1E; Figure S9). Interestingly, there are three distinct segments, primarily between residue 900 and the beginning of the Rap-GAP domain, where very few predicted classifications are above the pathogenic threshold. The Rap-GAP domain itself appears to be enriched for predicted pathogenic variants. We then investigated the overall distribution of TRUST’s output by class (Figure 1F). As expected, BLB variants have a low overall probability of pathogenicity, while PLP variants have a high probability of pathogenicity. Interestingly, the missense VUS population has a distribution that is most similar to the BLB class, suggesting the majority of VUS may be rare, likely benign variants. It is important to note, though, that a small but appreciable proportion of VUS are indeed predicted to be pathogenic, suggesting it is important to find methods to distinguish these needles from the proverbial haystack. To ask whether TRUST’s predictions, or those of the globally-trained VEPs, also translate accurately to VUS, we turned our attention to building a high-throughput assay for functional characterization of *TSC2* variants.

### Curated loci prime editing (cliPE)-mediated MAVE enables TSC2 VUS resolution

We previously developed a CRISPR/Cas9-induced homology directed repair approach to perform medium-throughput VUS resolution for about a dozen variants in the mTORopathy gene *SZT2*.^15^ Scaling this approach to perform VUS resolution on more than 100 *TSC2* missense VUS required modifications to the experimental workflow (Figure 2A). Primarily, we modified the approach to incorporate high-throughput prime editing of the endogenous *TSC2* locus of HAP1 cells. The prime editing portion of this workflow took inspiration from a recent study which performed saturation prime editing (SPE) on portions of two genes.^33^ SPE requires a pegRNA library containing every possible codon within an approximately 10-codon stretch, the estimated distance limit of efficient prime editing. To reduce cost, we focused on generating libraries enriched in missense VUS as well as control variants necessary for assay validation and calibration, including synonymous and missense variants present in a population database (gnomAD), ClinVar known missense BLB and PLP variants, and synthetic premature truncation codons (PTCs) to induce TSC2 LOF. With these libraries, HAP1 cells are co-transfected with prime editing reagents to generate up to ∼60 variants per target region, followed by serum starvation, FACS sorting for P-S6 (mTORC1 activity biomarker), targeted amplicon sequencing, and allele enrichment analysis.

**Figure 2:**
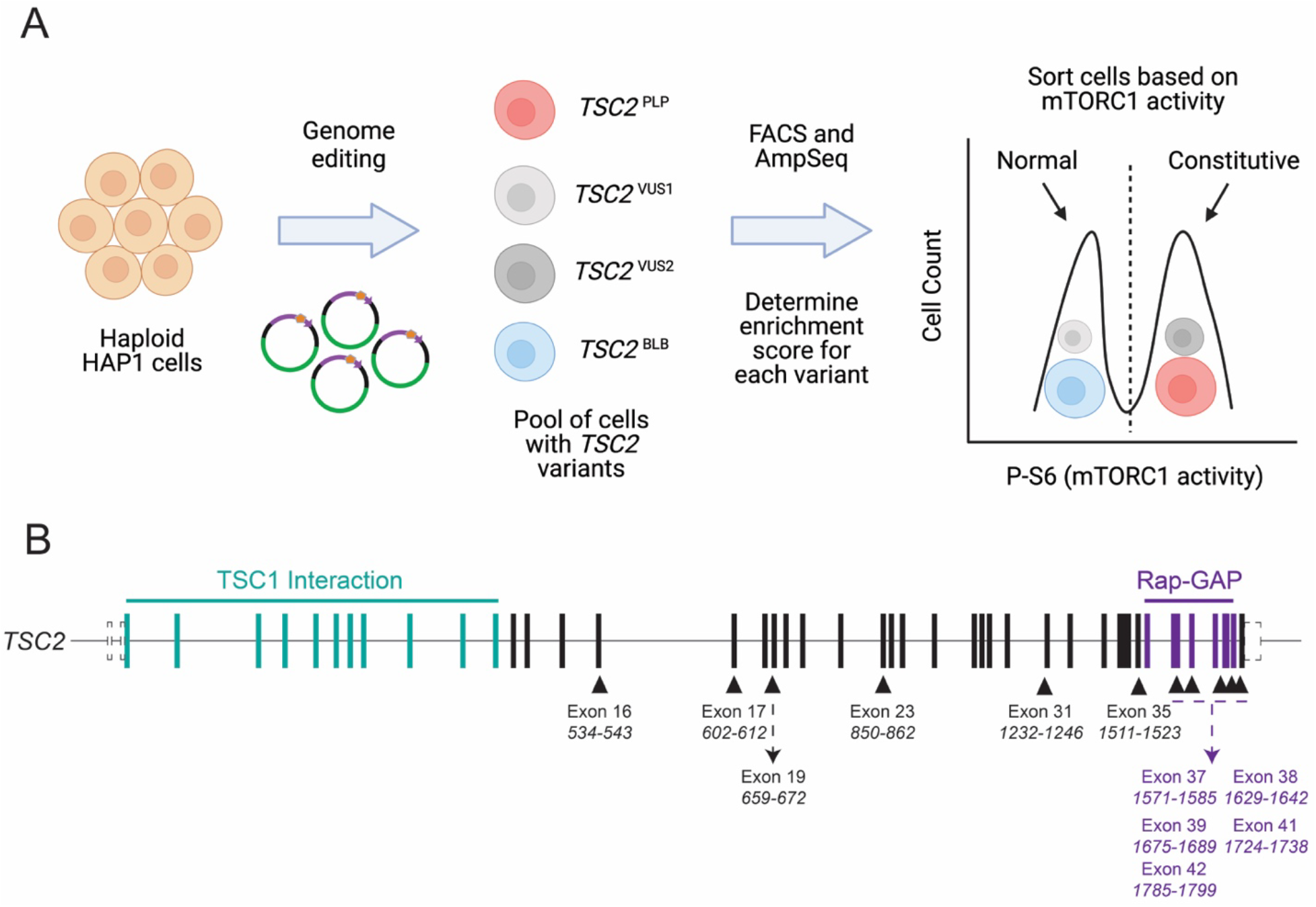
Overview of curated loci prime editing (cliPE) method for *TSC2* MAVE. (**A**) Cartoon overview showing a simplified version of the cliPE method for performing the *TSC2* MAVE reported herein. (**B**) epegRNA screen design based on regions of high local VUS density in *TSC2*. Exons targeted are shown and the codons overlapped by the epegRNA are provided. Codon numbering is based on *TSC2* transcript NM_000548.5.

We first screened 11 epegRNA architectures for prime editing efficiency as recommended by Doman JL et al. (Figure 2B).^25^ 6/11 (54.5%) exhibited robust prime editing. We then designed cliPE libraries for these 6 regions which could theoretically generate up to 300 *TSC2* variants. Our NGS amplicon sequencing in unsorted cells revealed that 69.4% of variants were successfully made in HAP1s (213/307; Table S5). We then performed enrichment analysis to determine CMAS scores for these 213 variants (Figure 3A; Figures S10 to S15). The CMAS distributions for synonymous and PTC variants are well-separated, suggesting the variants comprise distinct populations (Figure 3B). We determined thresholds for assigning pathogenicity based on the 95% confidence intervals for these two populations (T_BLB_ = 0.242; T_PLP_ = 0.477). Using these thresholds, we next investigated pathogenicity of known missense BLB and PLP variants. Of 25 total variants (13x BLB; 12x PLP), 23 were correctly classified with 2 variants remaining ambiguous (Figure 3C). This level of evidence is sufficient to ascribe moderate evidence of benignity or pathogenicity based on a Brnich et al. framework.^34^ We generated functional scores for 106 *TSC2* VUS. Though we do not have precise clinical information on individuals with VUS in ClinVar, we can still hypothetically perform VUS resolution under some relatively safe assumptions using the ACMG framework.^35^ First, *in silico* prediction of benignity (supporting; BP4) and MAVE evidence of benignity (moderate; BS3) will be sufficient to reclassify VUS to likely benign in many instances. Secondly, *in silico* prediction of pathogenicity (supporting; PP3), MAVE evidence of pathogenicity (moderate; PS3), *de novo* or assumed (moderate PM6 or strong PS2), and absence from controls (moderate; PM2) will be sufficient to reclassify VUS to likely pathogenic in some instances. Under these assumptions, we are able to resolve 92 unique VUS from ClinVar; 86 variants were resolved to likely benign and 6 variants were resolved to likely pathogenic (Figure 4).

**Figure 3:**
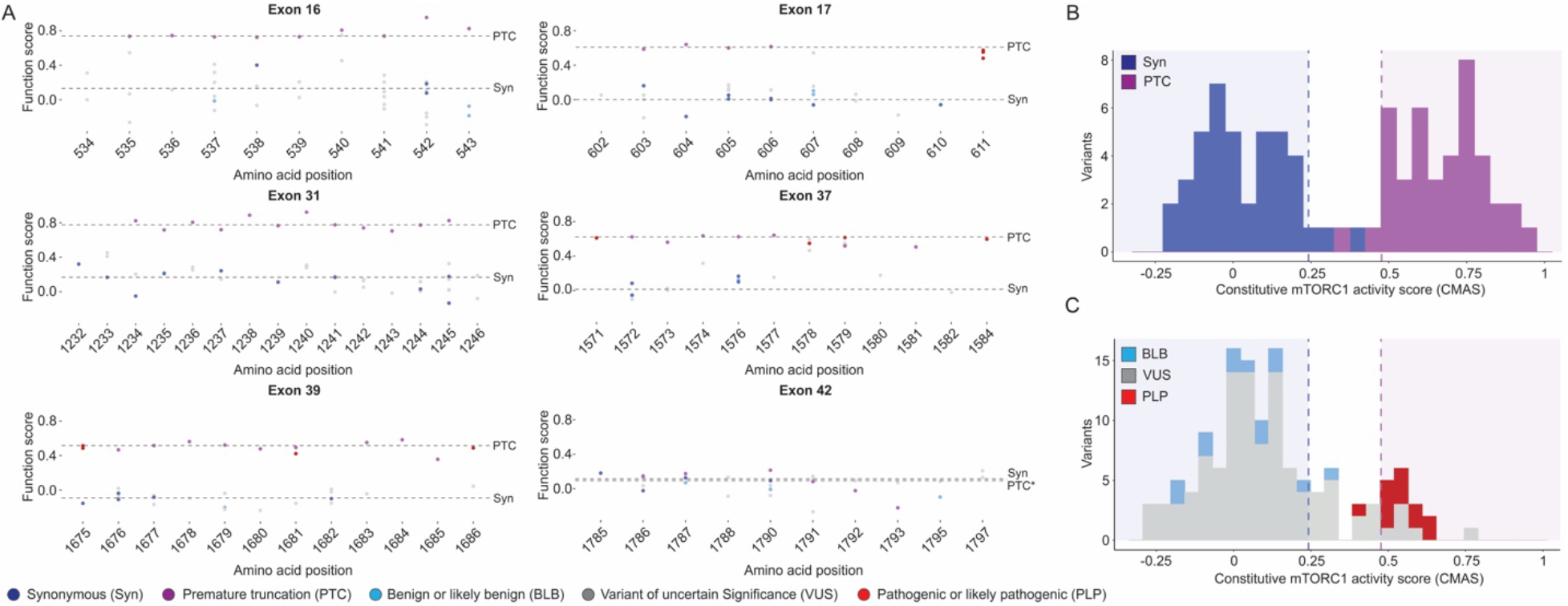
cliPE mapping of variant effect in more than 200 variants, including 106 missense VUS. (**A**) CMAS scores plotted by amino acid position and grouped by epegRNA target region. Scores are derived from 3-4 biological replicates. Dashed lines indicate the median score for either synonymous (Syn) or PTC variants for that particular exon. Plots of each variant with standard error are provided in Figures S10 to S15. *Exon 42 is the final exon in TSC2 and as such PTC variants likely escape nonsense-mediated decay and encode functional proteins, as evidenced by the median score for synonymous and PTC variants being equivalent. (**B & C**) Histograms of CMAS, bars colored by variant type. Synonymous (n=41) and PTC (n=41) variants (excluding final exon 42 PTCs) used to determine pathogenicity thresholds are shown in the top plot (**B**). Missense BLB (n=13), PLP (n=12), and VUS (n=106) are shown in the bottom plot (**C**). Full details of the truth set can be found in Table S4.

**Figure 4:**
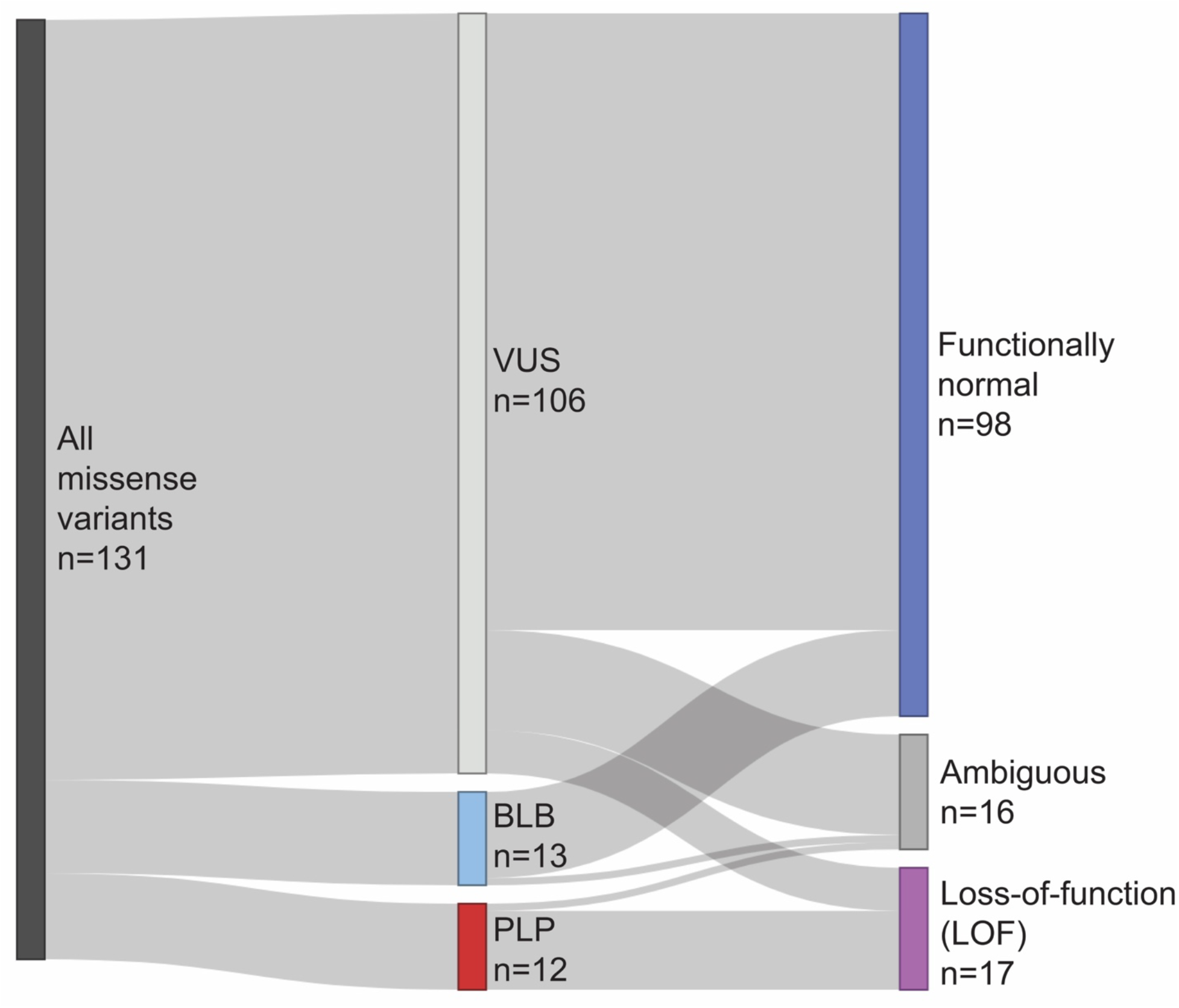
*TSC2* MAVE has the potential to reclassify many missense VUS. Sankey diagram showing that with moderate strength evidence of benignity or pathogenicity, the current *TSC2* MAVE data could enable reclassification of up to 86.8% of missense VUS. In agreement with our *in silico* classifier, most *TSC2* missense VUS appear to be rare benign variants.

### TRUST predictions align with TSC2 MAVE scores

The missense variants investigated in our MAVE constitute an independent holdout dataset with which to evaluate the performance of TRUST. We first asked how well our pathogenicity classifications from TRUST agreed with the pathogenicity determinations from the MAVE. TRUST’s predictions agreed well with the MAVE classifications (96.9% specificity; 95.6% accuracy) (Figure 5A-B; Figure S16; Table 3). Furthermore, the probability of pathogenicity scores from TRUST correlate well with the CMAS scores derived from the MAVE (Figure 5C; Figures S17 and S18). Based on our evidence across a truth set and two independent holdout datasets, we posit TRUST has utility for predicting pathogenicity for variants where we currently lack functional characterization (Figure 5D).

**Table 3:** Machine learning model metrics on *TSC2* MAVE dataset.

| Model | Threshold | Specificity | Recall | Accuracy | Precision | F1 score | Spearman correlation with CMAS | Pearson correlation with CMAS |
| --- | --- | --- | --- | --- | --- | --- | --- | --- |
| TRUST | 0.4684 | 0.9691 | 0.8824 | 0.9561 | 0.8333 | 0.8571 | 0.3862 | 0.6040 |
| Alpha Missense | 0.8153 | 0.9588 | 0.8824 | 0.9474 | 0.7895 | 0.8333 | 0.5452 | 0.6078 |
| BayesDel | 0.3428 | 0.8866 | 0.8824 | 0.8860 | 0.5769 | 0.6977 | 0.3730 | 0.4542 |
| ClinPred | 0.9456 | 0.7835 | 0.8824 | 0.7982 | 0.4167 | 0.5660 | 0.3850 | 0.2944 |
| EVE | 0.7068 | 0.8889 | 0.8235 | 0.8776 | 0.6090 | 0.7000 | 0.3955 | 0.4691 |
| gMVP | 0.7490 | 0.9691 | 0.8824 | 0.9561 | 0.8333 | 0.8571 | 0.4143 | 0.4984 |
| REVEL | 0.7880 | 0.9485 | 0.8235 | 0.9298 | 0.7368 | 0.7778 | 0.2816 | 0.3834 |
| VARITY_R | 0.7563 | 0.9485 | 0.8235 | 0.9298 | 0.7368 | 0.7778 | 0.3345 | 0.4545 |

**Figure 5:**
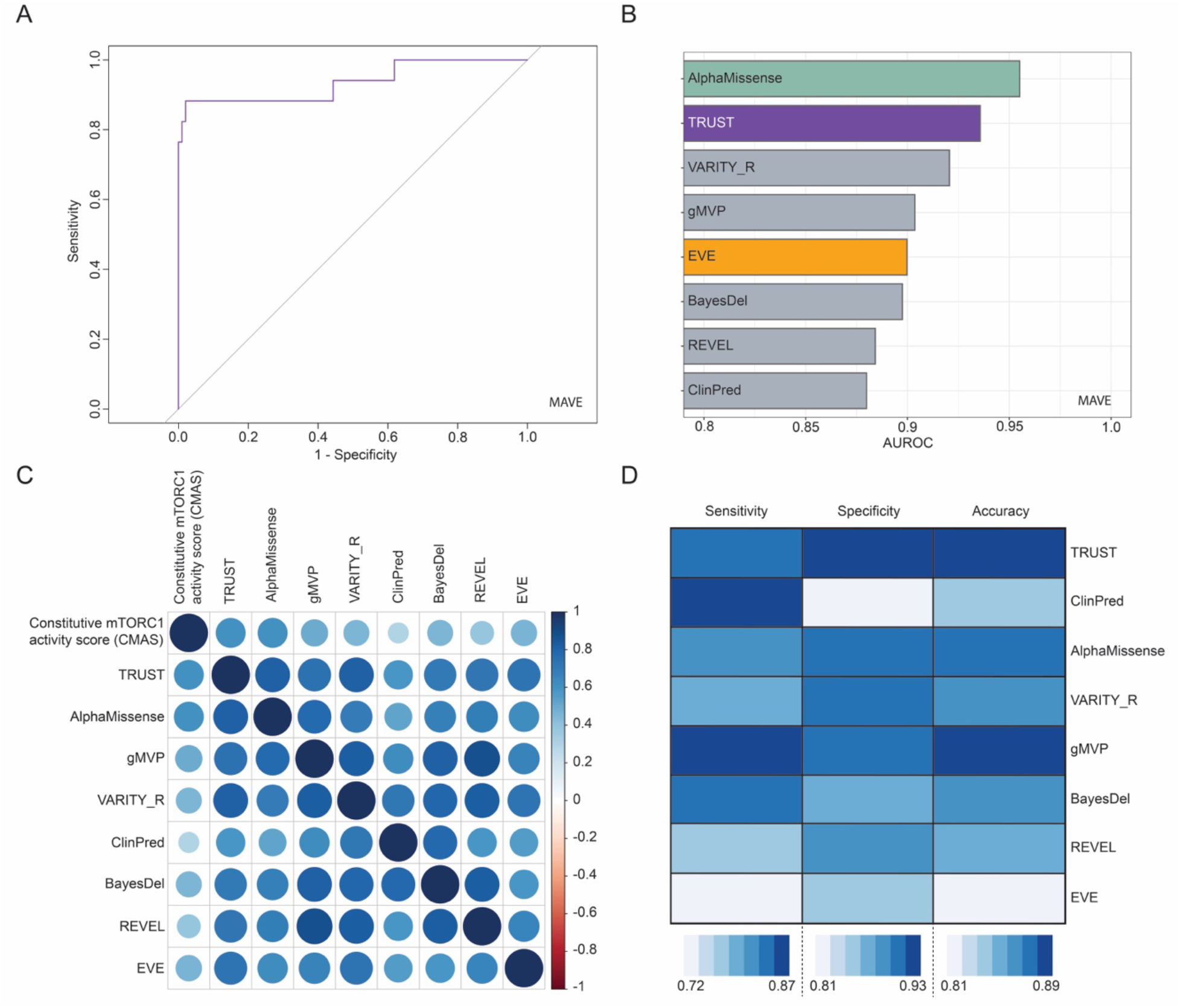
Validation of gene-specific VEP performance on MAVE data. (**A**) Receiver operator characteristic (ROC) curve for TRUST classifier on missense variants classified unambiguously by CMAS score from the *TSC2* MAVE (n=114 variants). (**B**) Area under the ROC curve (AUROC) for TRUST and some of the best-performing globally-trained VEPs on the MAVE dataset. (**C**) Correlation (Pearson) plot of TRUST, CMAS, and some of the best-performing globally-trained VEPs. (**D**) Heatmap of overall performance of gene-specific *in silico* classifier and best-performing globally-trained VEPs averaged across all three classification datasets (ClinVar test set, holdout set, and MAVE data).

### Global explanations reveal protein stability as a possible molecular phenotype for TSC2 missense variants

One of the limitations of supervised learning models is their “black box” nature; it can be difficult to impossible to understand the logic behind a model’s predictions. A recent method, Shapley additive explanations (SHAP), uses game theory to investigate a classifier’s predictions.^36^ We used scikit-learn’s feature importance and SHAP’s kernel explainer to investigate TRUST’s global explanations, or the features most contributing to its final predictions. Both tools agreed on the top 10 features, which included other global VEPs (AlphaMissense, PROVEAN, etc.), relative solvent accessibility, and MAESTRO, a predictor of protein stability (Figure 6). Interestingly, all three (iMutant, mCSM, and MAESTRO) *in silico* protein stability predictor features are among the top features, suggesting protein stability may be a molecular mechanism worth investigating in more detail.

**Figure 6:**
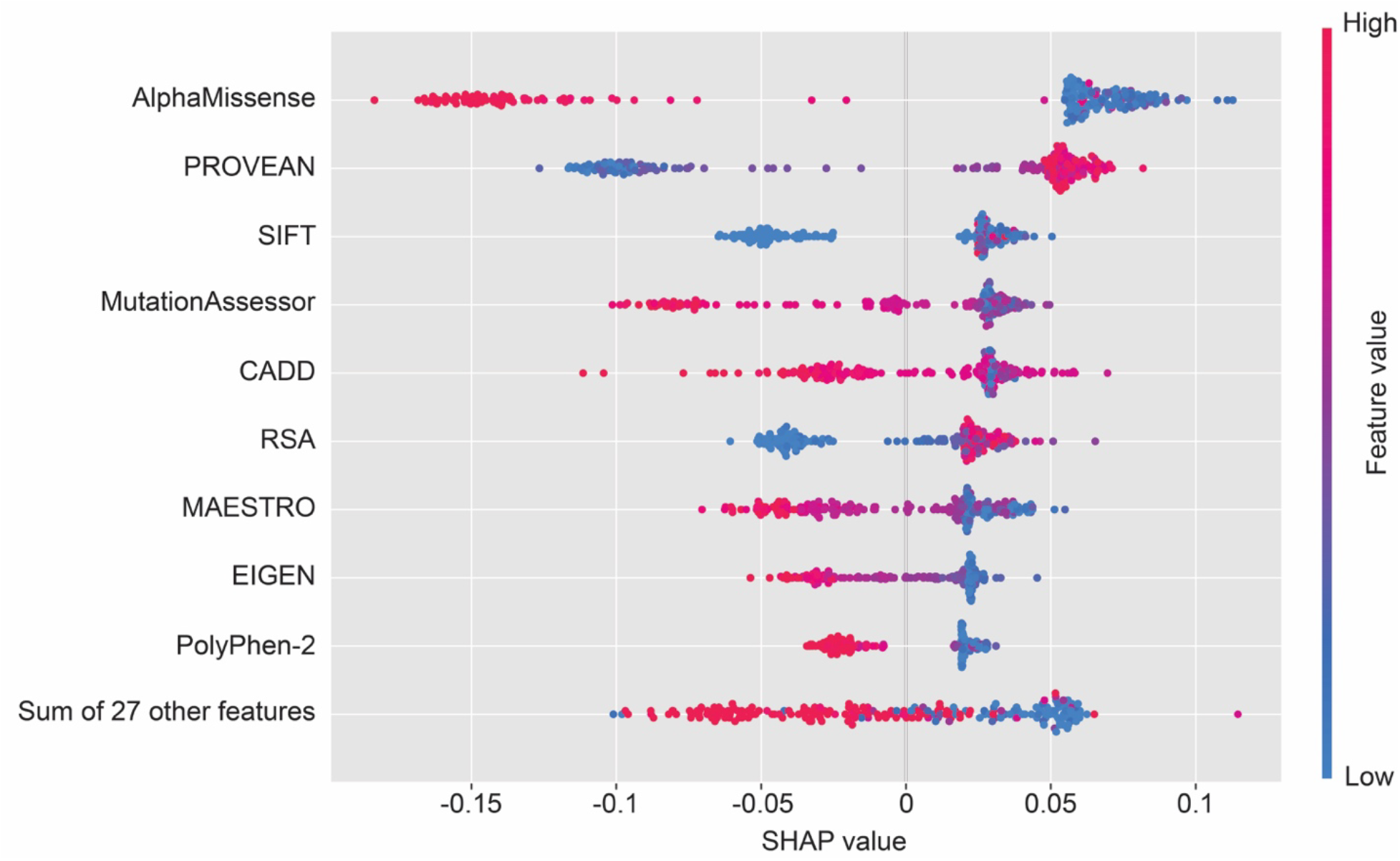
Global explanations provide insight into *TSC2* gene-specific *in silico* pathogenicity prediction. Contributions of features to final model as determined by SHAP values. Other globally-trained VEPs including AlphaMissense, PROVEAN, SIFT, and PolyPhen-2 are among the top features. Notably, a structure-related feature (relative solvent accessibility; RSA) and an *in silico* predictor of protein stability (MAESTRO) are also among the top features, suggesting that missense variants in *TSC2* may impact protein folding or abundance.

## Discussion

Herein, we report TRUST, a new gene-specific VEP for *in silico* pathogenicity prediction of missense *TSC2* variants. This algorithm is a random forest trained on more than 300 known BLB and PLP missense variants from the ClinVar database. We used an independent holdout dataset of missense variants to demonstrate that TRUST’s predictions generalize well. Overall, TRUST outperforms globally-trained modern VEPs such as AlphaMissense, VARITY, or REVEL across the full truth set and our two holdout datasets. It is worth noting that there are a few instances where a globally-trained VEP outperforms TRUST, such as gMVP or ClinPred on the test dataset. We cannot rule out a possible impact of type 1 circularity for many of these globally-trained VEPs; the variants in our test dataset were likely present in the respective training sets of these VEPs. However, it is difficult to gauge the impact of this circularity, given these VEPs have large, genome-wide training sets. We provide predictions for all possible SNVs in *TSC2* coding exons which produce a missense variant (Table S2). We posit that these predictions may have utility as supporting evidence for benignity or pathogenicity within an ACMG variant classification framework.

We also developed cliPE, which allowed us to generate over 200 variants across 6 exons of *TSC2*. We adapted our previous *SZT2* VUS resolution assay and combined it with cliPE to generate functional scores (CMAS) for synonymous, missense, and PTC variants. PTC variants, but not synonymous variants, were enriched in cells with high mTORC1 activity. Missense PLP variants, but not BLB variants, were similarly enriched in cells with constitutive mTORC1 activity. Our MAVE data allowed resolution of 92/106 VUS, with the majority of variants resolved to BLB. This agrees with TRUST’s prediction that about 78% (8,300/10,623) of scored missense variants are likely to retain function.

This study has two limitations that are worth noting. First, there is a tradeoff in cost between cliPE and saturation genome editing (SGE) or SPE. SGE and SPE are more expensive in terms of experimenter time, reagent cost, and sequencing cost. However, SGE/SPE generate all possible missense variants in a targeted region. As such, those methods are both retrospective and prospective; they generate MAVE data for variants that have and have yet to be sequenced in an individual. cliPE is purely retrospective, and its variant generation is dependent on databases such as ClinVar, gnomAD, the Regeneron million exome server, etc. We recommend cliPE as a good alternative to SPE when budget is a limiting factor; SGE/SPE should be considered if funding allows.

Second, low-passage HAP1 cells were used in this study, but we did not select for ploidy using flow cytometry or pharmacology.^37, 38^ As *TSC2* is an autosomal dominant disorder, the ploidy of cells is unlikely to be a major confounding factor. However, we cannot rule out a small effect of ploidy on our assay results, particularly for putative partial LOF variants or dominant negative variants. Our primary goal was developing a MAVE which could distinguish complete LOF missense variants from variants retaining function. Future work will be necessary to develop assays that are calibrated to test for partial LOF or dominant negative variants.

This study builds on the excellent work by the Nellist lab and others to functionally characterize individual missense variants in *TSC2*.^39^ We have made TRUST’s predictions and MAVE data available in the supplementary information (Table S2 and Tables S4-5). Our plan is to build a website where we can more readily share this data in a format that is accessible to clinicians, genetic counselors, and the TSC community. Our MAVE is registered on the MAVE Registry, and we will deposit our data in MAVEdb.^40, 41^ We also view this study in many ways as a first step towards comprehensive VUS resolution in *TSC2* specifically, and the mTORopathies more broadly. The methods utilized herein are generalizable and could be used to study mTORopathy genes with hundreds of missense VUS such as *MTOR* and *PIK3CA*, perhaps increasing access to mTOR inhibitors and other forms of precise clinical management. cliPE also may have broader utility to enable accessible MAVEs in other genes beyond the mTORopathies.

## Supporting information

Supplemental Figures and References

## Web resources

gnomAD v4 (https://gnomad.broadinstitute.org/); accessed 5 Jan 2024 for curation of holdout dataset

Regeneron Genetics Center (RGC) Million Exome Variant Browser (https://rgc-research.regeneron.com/me/license-and-terms-of-use); accessed 5 Jan 2024 for curation of holdout dataset

ClinVar (https://www.ncbi.nlm.nih.gov/clinvar/); accessed 24 Oct 2022 for initial list of BLB, PLP, and VUS for design of epegRNA libraries; accessed 18 Dec 2023 for final list of BLB, PLP, and VUS for supervised machine learning

*TSC2* LOVD gene-specific database (https://databases.lovd.nl/shared/genes/TSC2); accessed 8 Jan 2024 for curation of holdout dataset

Some figure panels were created with BioRender.com.

## Acknowledgments

This work was supported by the Northwestern University – Flow Cytometry Core Facility supported by Cancer Center Support Grant (NCI CA060553). Flow Cytometry Cell Sorting was performed on a BD FACSAria SORP system and BD FACSymphony S6 SORP system, purchased through the support of NIH 1S10OD011996-01 and 1S10OD026814-01. We thank David Liu and the Liu lab for sharing their prime editing reagents. The authors would like to acknowledge Addgene for its invaluable service which facilitated this study. We further thank the Plasmidsaurus sequencing team for making quality control and pilot experiments more feasible and accessible than in the past.

