## Supplemental Figures and References for "Multimodal framework to resolve variants of uncertain significance in *TSC2*"

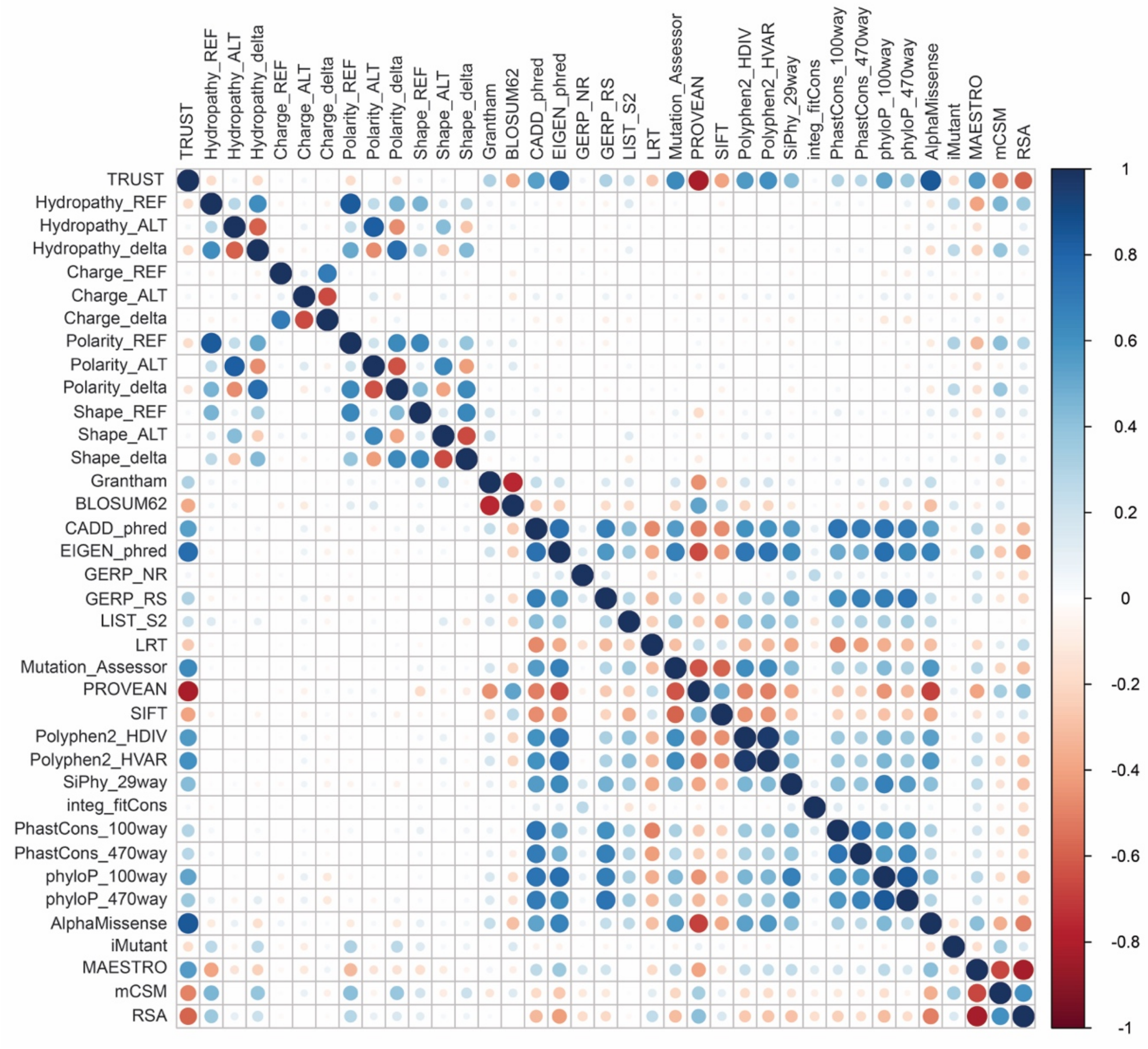

**Figure S1: Correlation plot of input features across full dataset.** Scores derived using Pearson correlation metric.

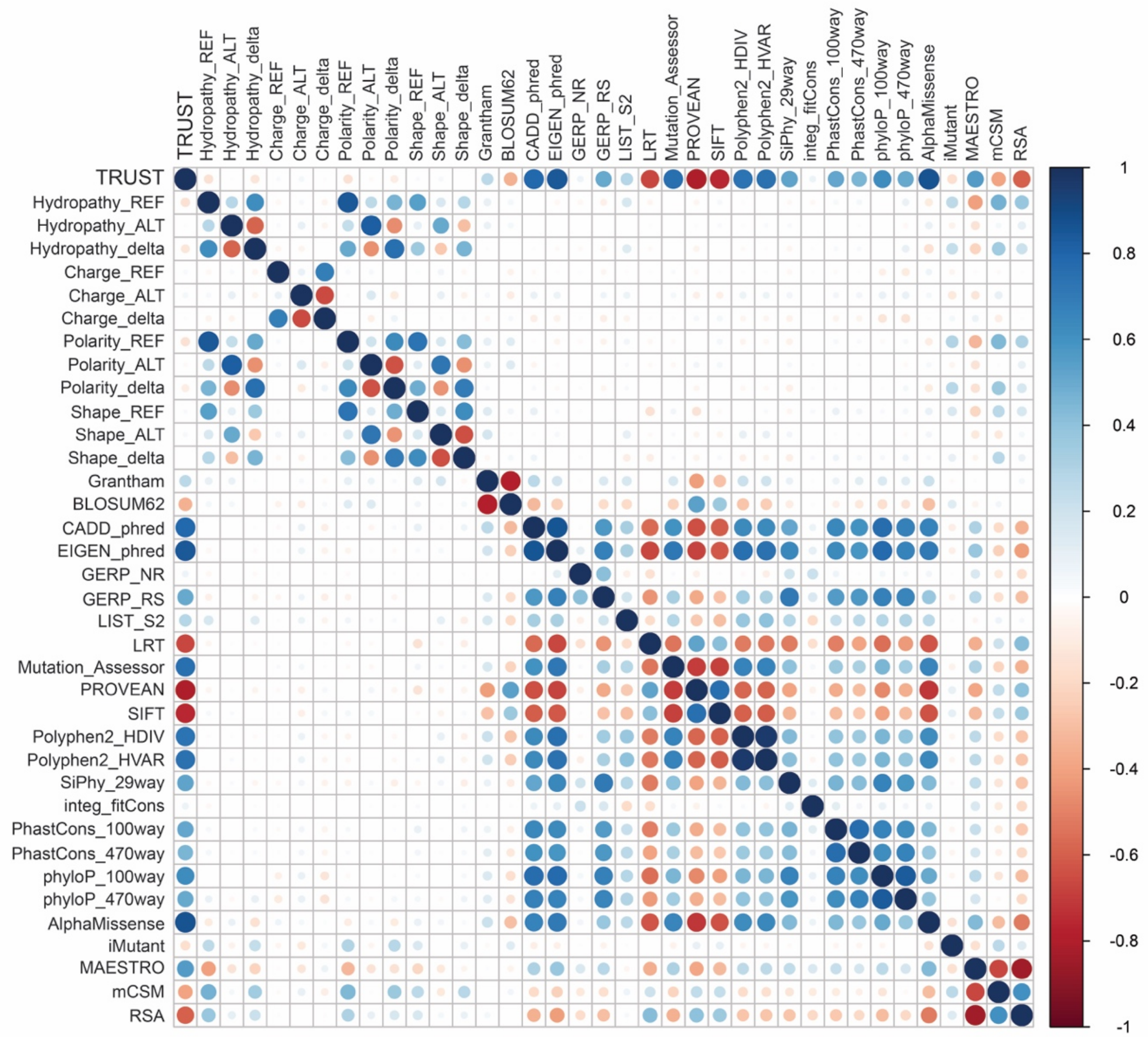

**Figure S2: Correlation plot of input features across full dataset.** Scores derived using Spearman correlation metric.

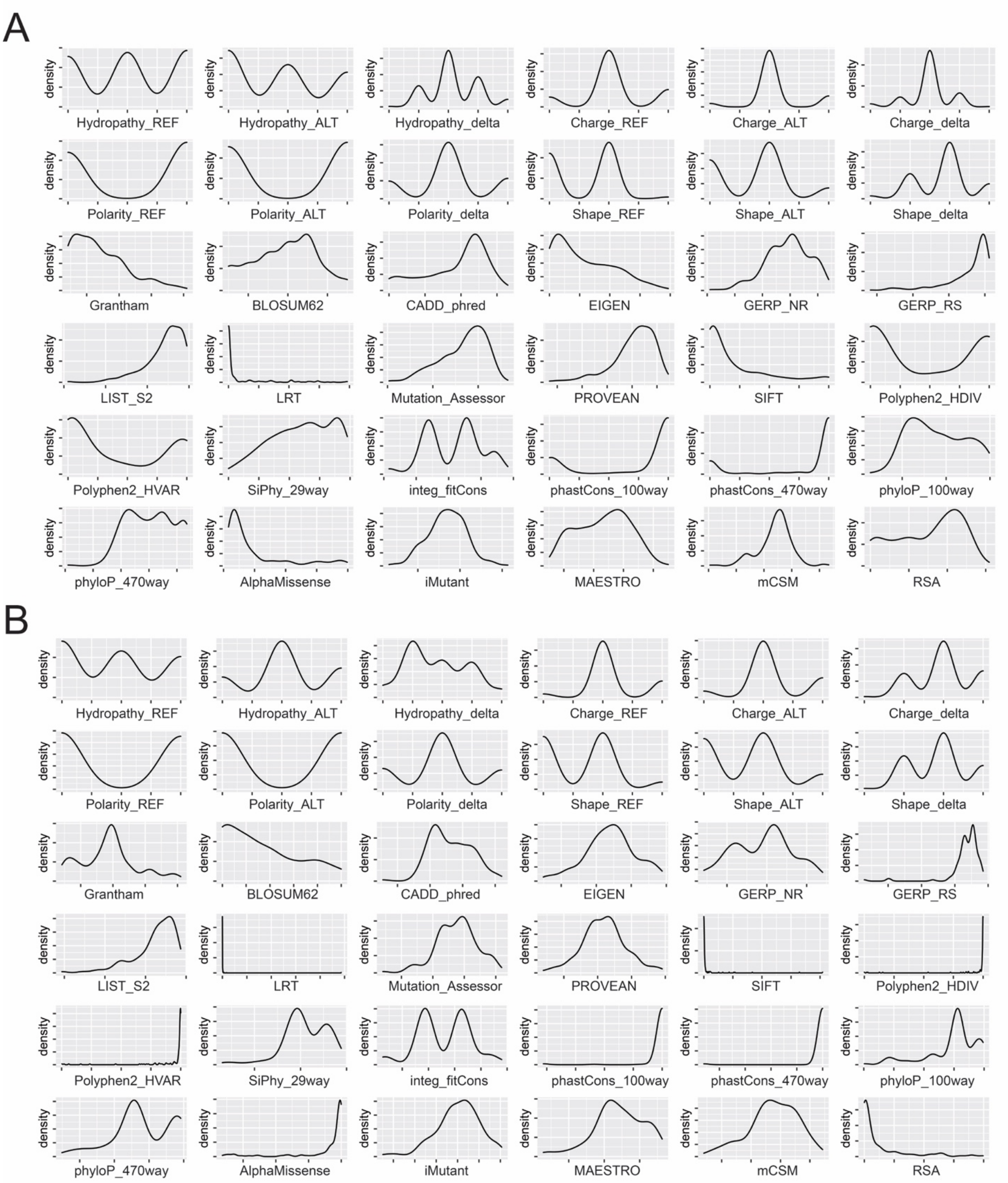

**Figure S3: Distributions of features used in model training.** (**A**) Histograms for each feature for class 0 variants (truth set BLB from ClinVar). (**B**) Histograms for each feature for class 1 variants (truth set PLP from ClinVar).

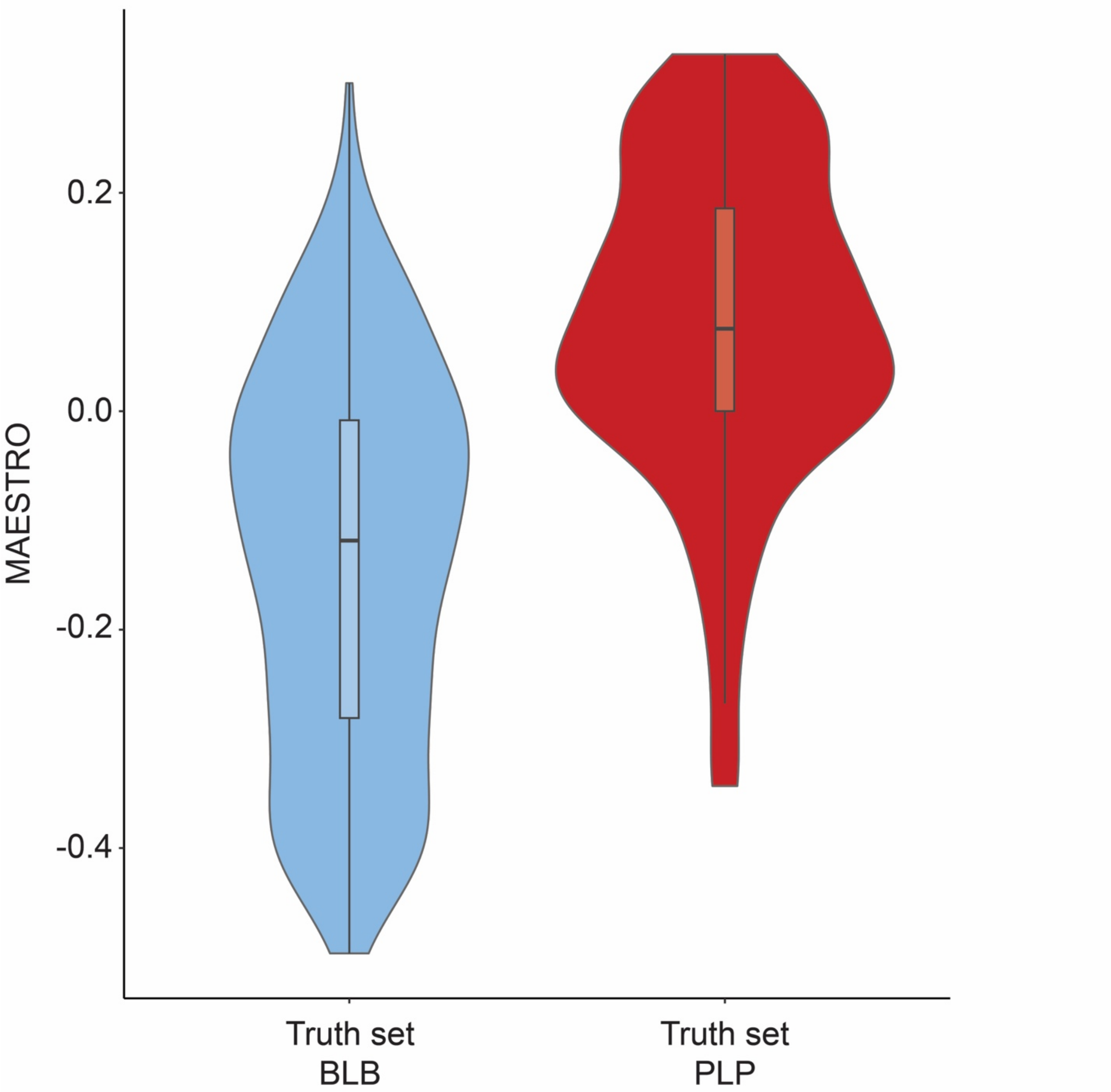

**Figure S4: Distributions of MAESTRO *in silico* stability predictions used in model training.** Violin plots for class 0 variants (truth set BLB from ClinVar) and class 1 variants (truth set PLP from ClinVar). $p < 0.001$ (Welch's t-test).

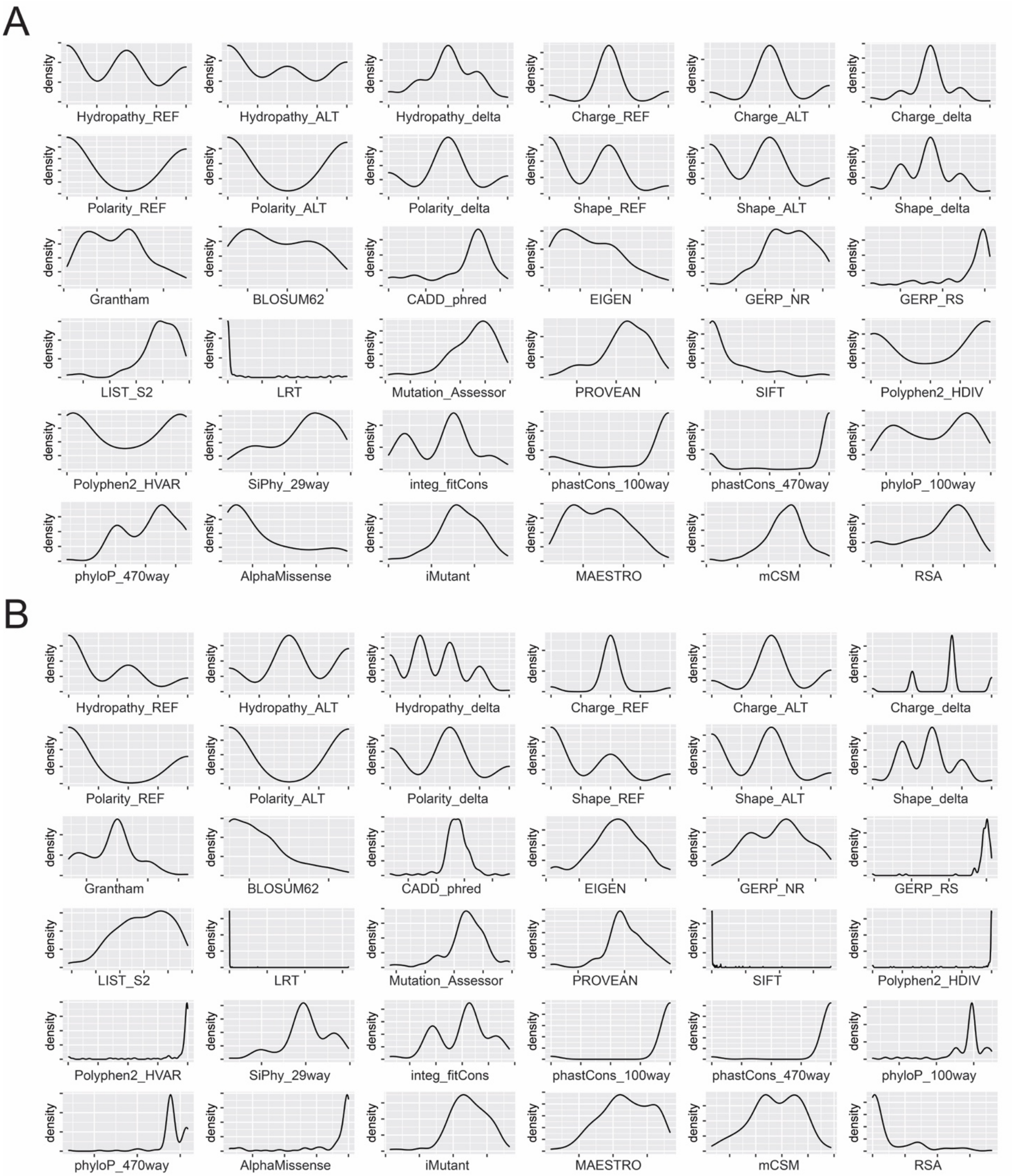

**Figure S5: Distributions of features in holdout dataset.** (**A**) Histograms for each feature for class 5 variants (holdout set BLB). (**B**) Histograms for each feature for class 6 variants (holdout set PLP).

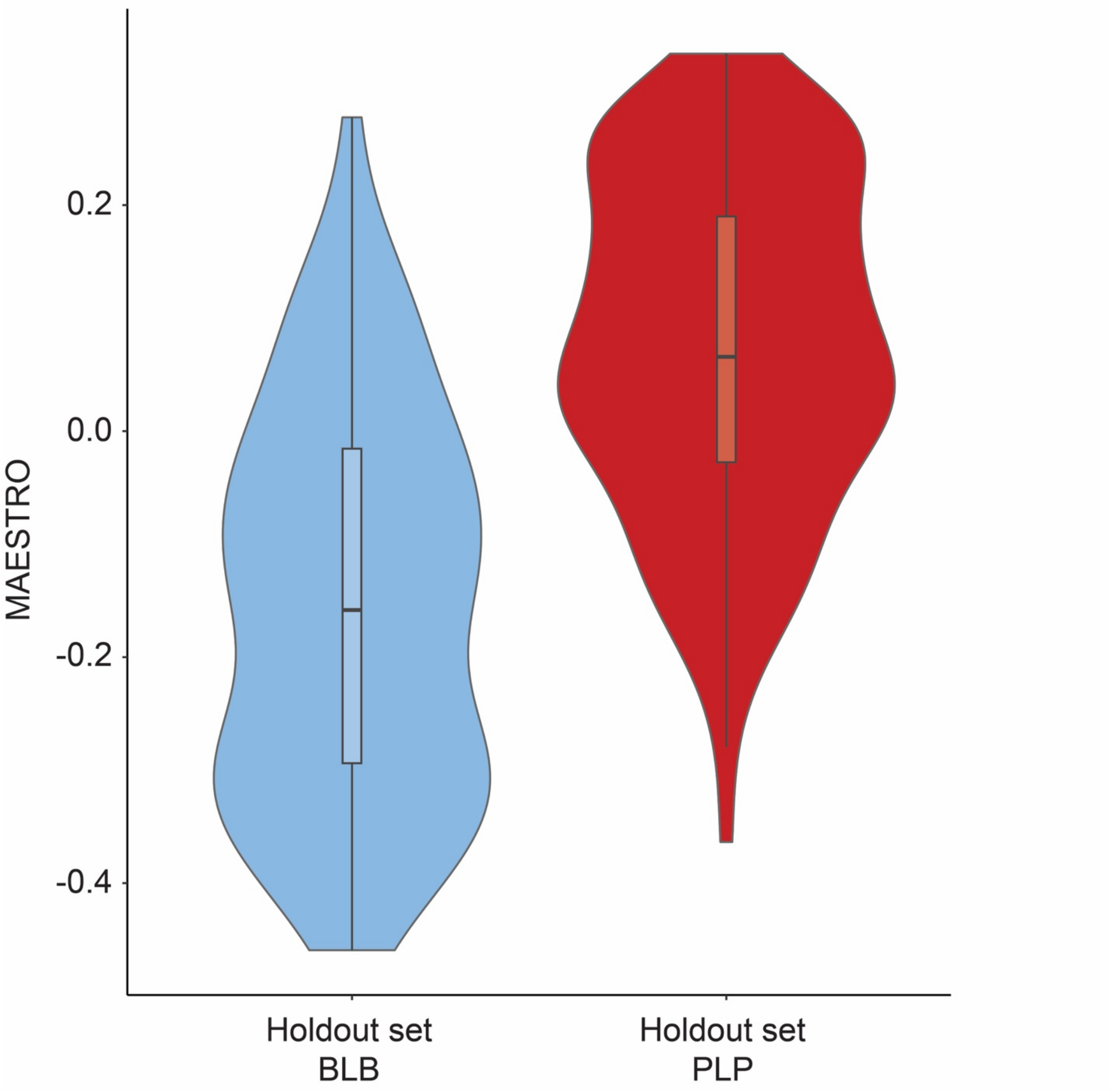

**Figure S6: Distributions of MAESTRO *in silico* stability predictions in holdout dataset.** Violin plots for class 5 variants (holdout set BLB) and class 6 variants (holdout set PLP). $p < 0.001$ (Welch's t-test).

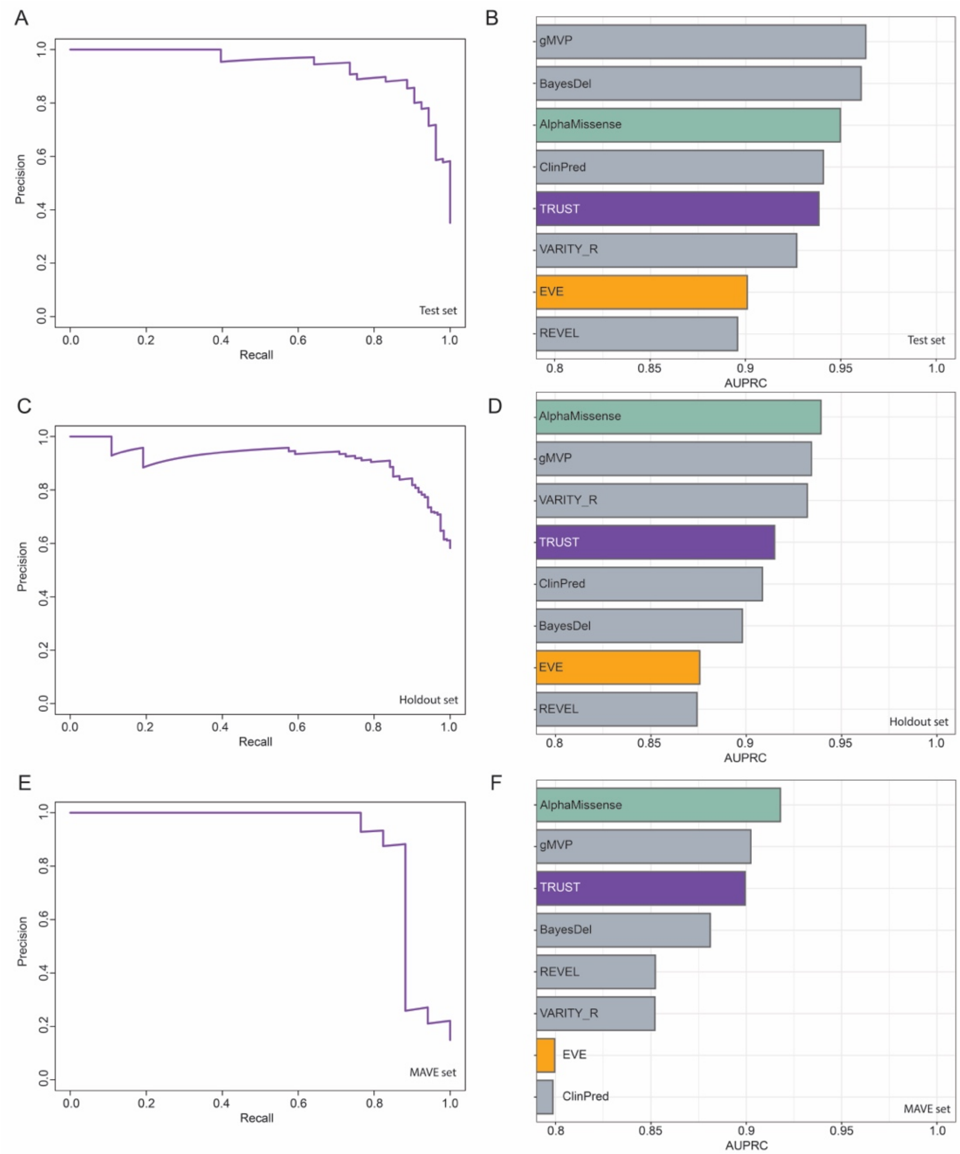

**Figure S7: Further evaluation of TRUST for predicting pathogenicity of missense variants.** (**A**) Precision recall (PR) curve for TRUST on the test set (n=151 variants). (**B**) Area under the PR curve (AUPRC) for TRUST and some of the best-performing globally-trained VEPs on the test set. Globally-trained VEPs trained on the ClinVar dataset are displayed in gray bars. (**C**) PR curve for TRUST classifier on the holdout dataset (n=206 variants). (**D**) AUPRC for TRUST and some of the best-performing globally-trained VEPs on the holdout dataset. (**E**) PR curve for TRUST on the MAVE dataset (n=114 variants). (**F**) AUPRC for TRUST and some of the best-performing globally-trained VEPs on the MAVE dataset.

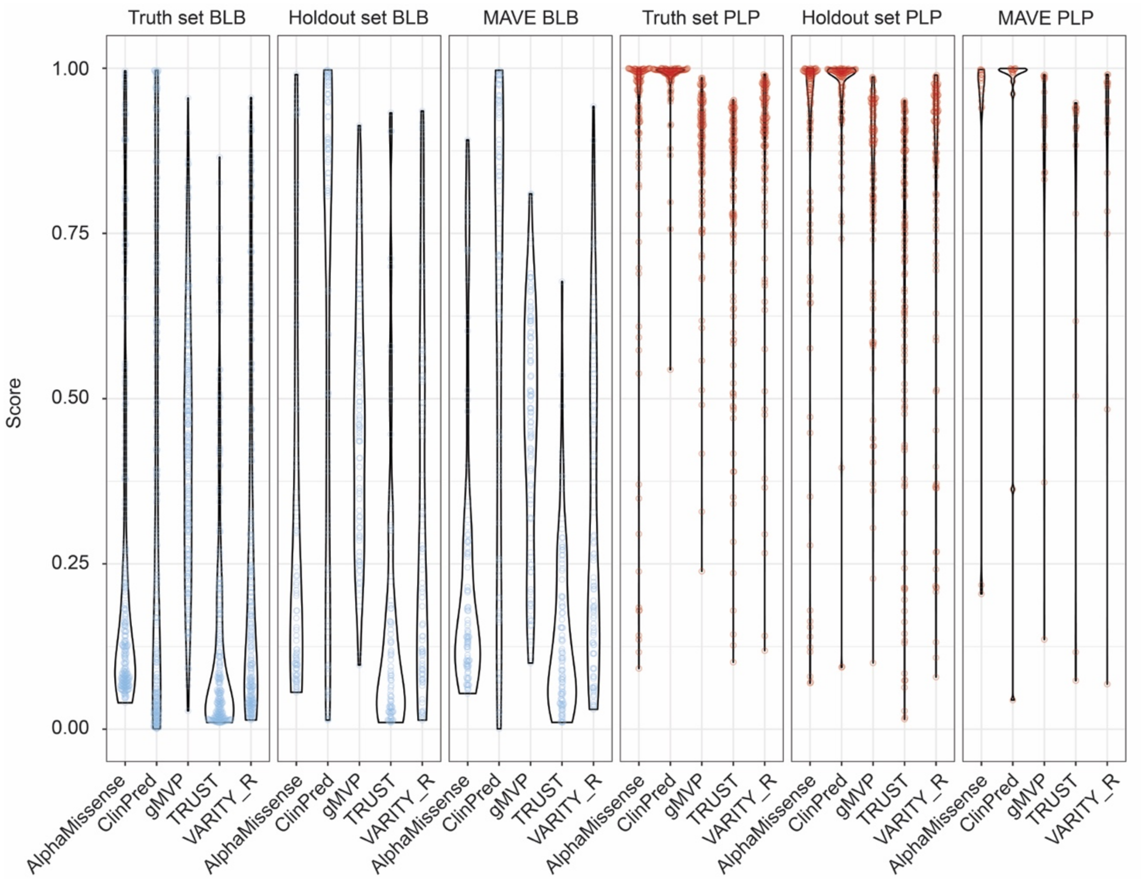

**Figure S8: Distributions of TRUST predictions and global VEP predictions across three datasets.** Violin plots of model outputs for each missense variant in truth set, holdout dataset, and MAVE dataset.

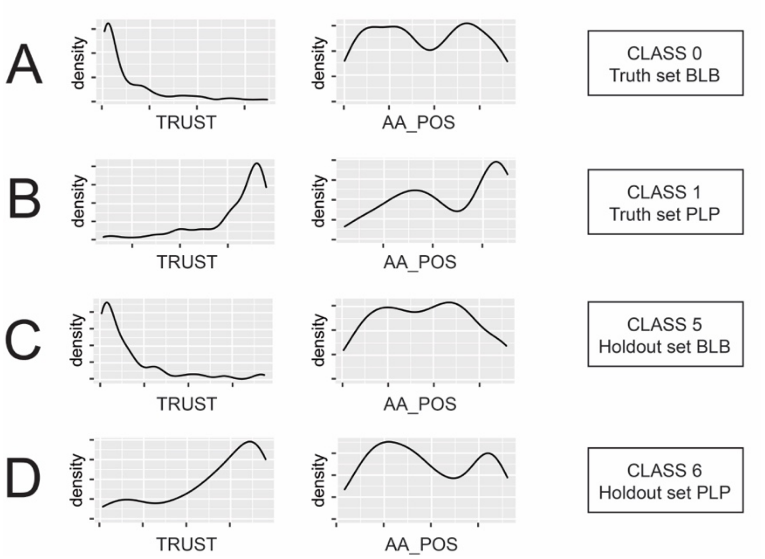

**Figure S9: Distributions of TRUST predictions and amino acid position in truth set and holdout dataset.** Histograms of probability of pathogenicity (TRUST) and amino acid position (AA_POS) for: (**A**) class 0 variants (truth set BLB from ClinVar), (**B**) class 1 variants (truth set PLP from ClinVar), (**C**) class 5 variants (holdout set BLB), and (**D**) class 6 variants (holdout set PLP).

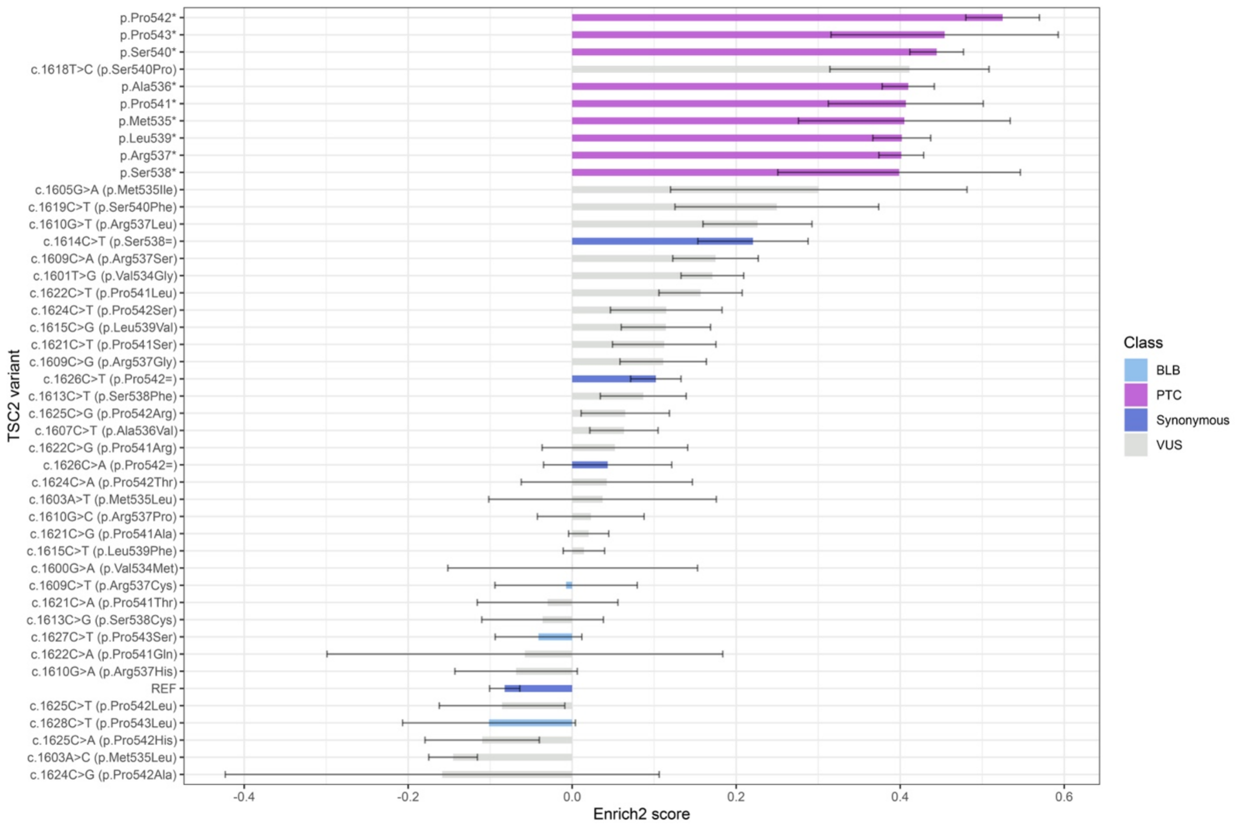

**Figure S10: Raw Enrich2 scores for each *TSC2* variant in exon 16.** Bars show combined enrichment score derived from 3 biological replicates. Error bars show standard error.

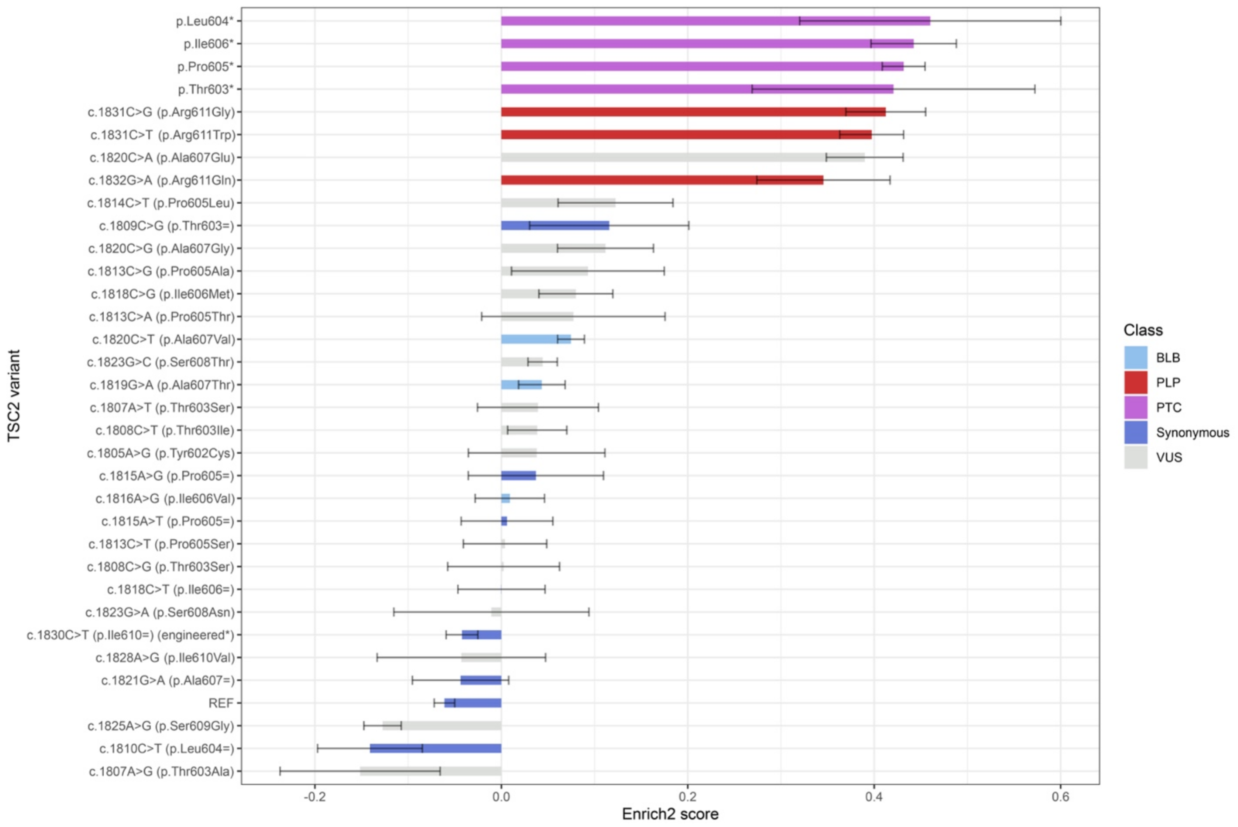

**Figure S11: Raw Enrich2 scores for each *TSC2* variant in exon 17.** Bars show combined enrichment score derived from 3-4 biological replicates. Error bars show standard error.

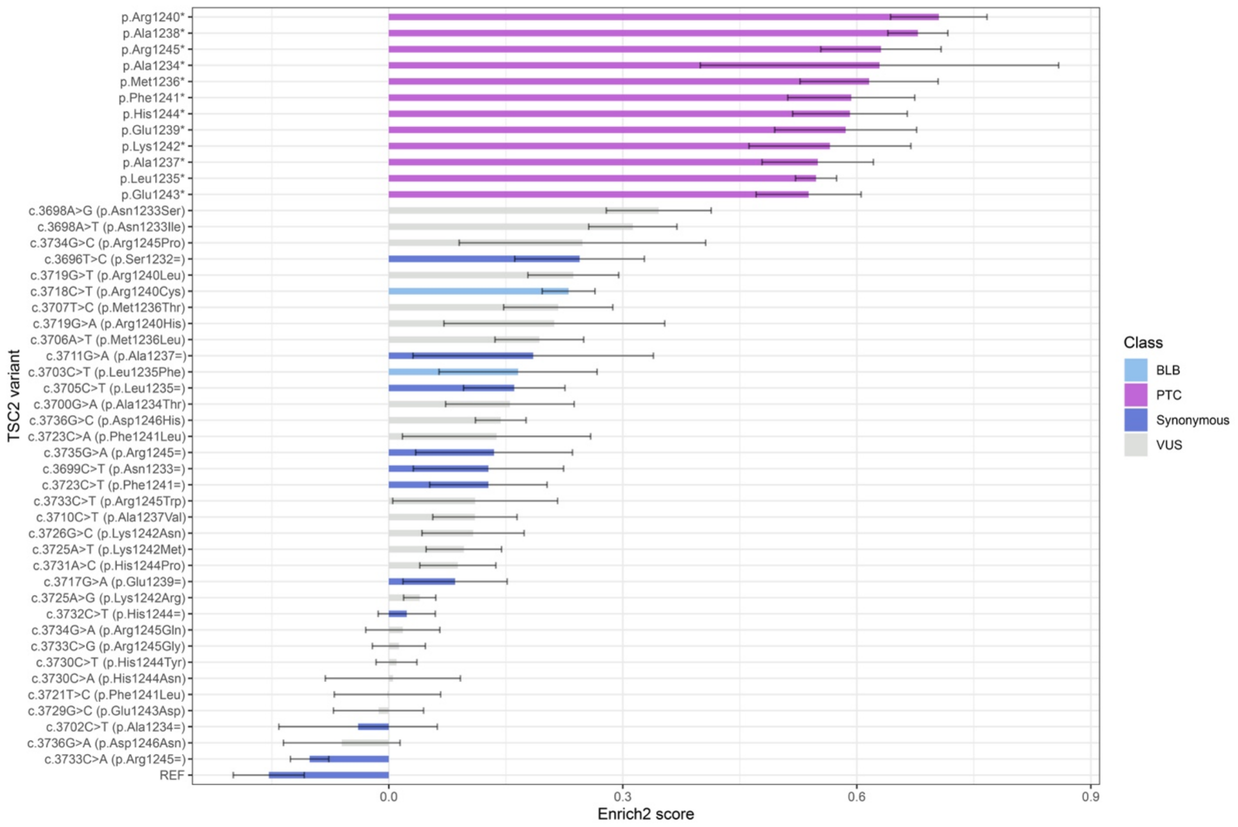

**Figure S12: Raw Enrich2 scores for each *TSC2* variant in exon 31.** Bars show combined enrichment score derived from 3-4 biological replicates. Error bars show standard error.

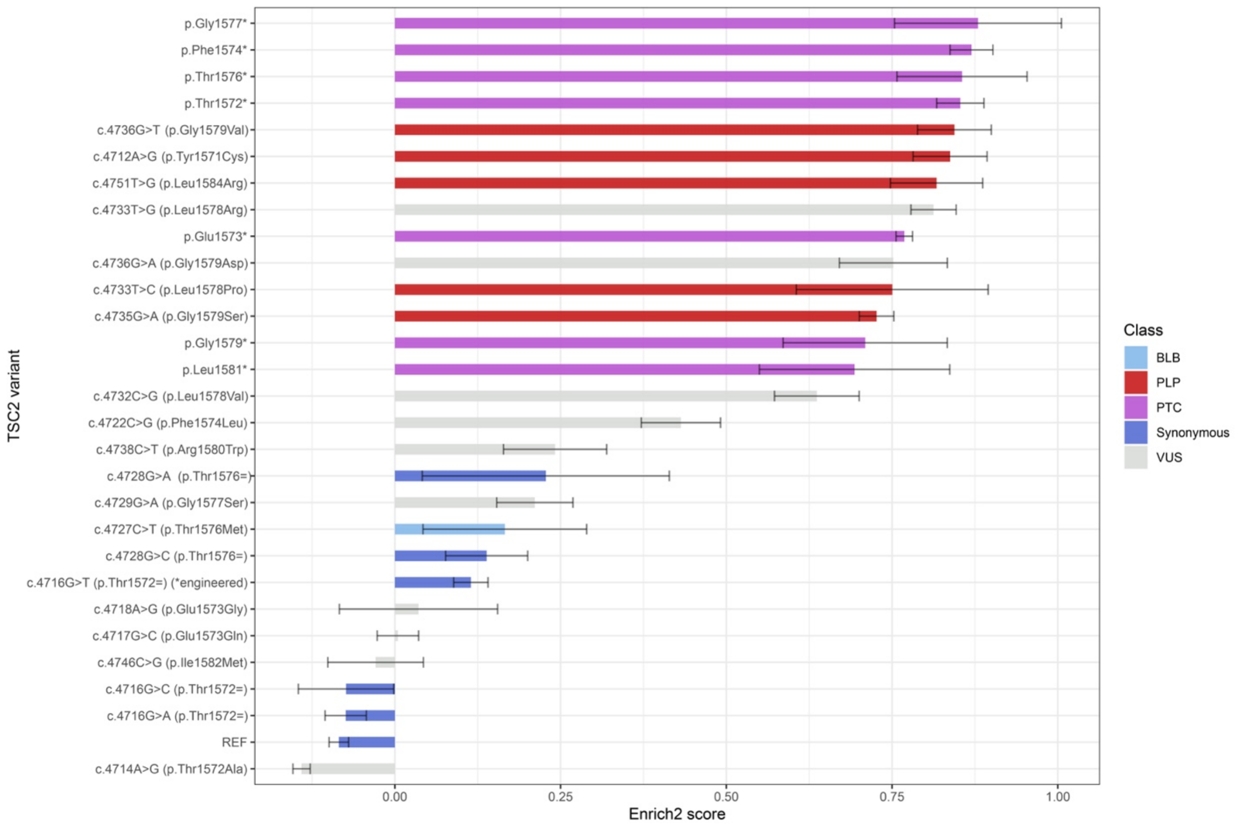

**Figure S13: Raw Enrich2 scores for each *TSC2* variant in exon 37.** Bars show combined enrichment score derived from 3-4 biological replicates. Error bars show standard error.

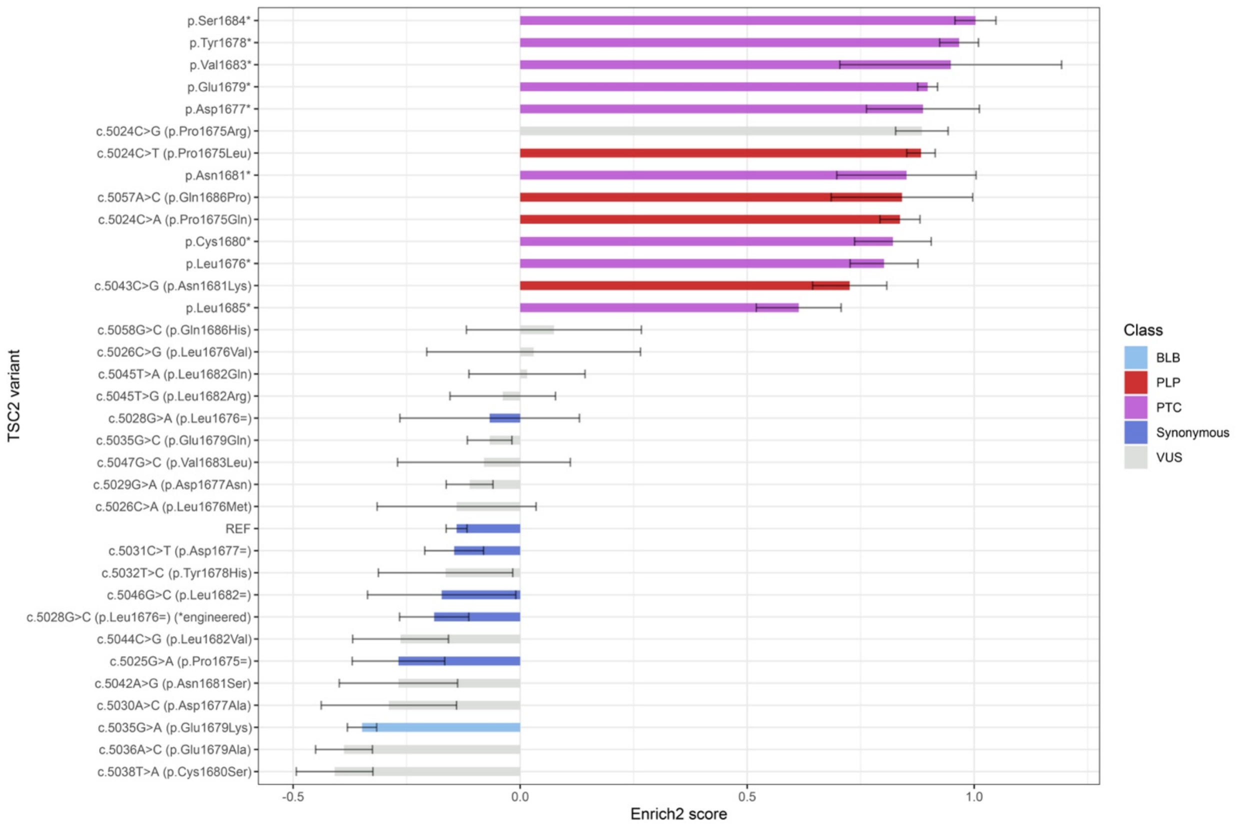

**Figure S14: Raw Enrich2 scores for each *TSC2* variant in exon 39.** Bars show combined enrichment score derived from 3 biological replicates. Error bars show standard error.

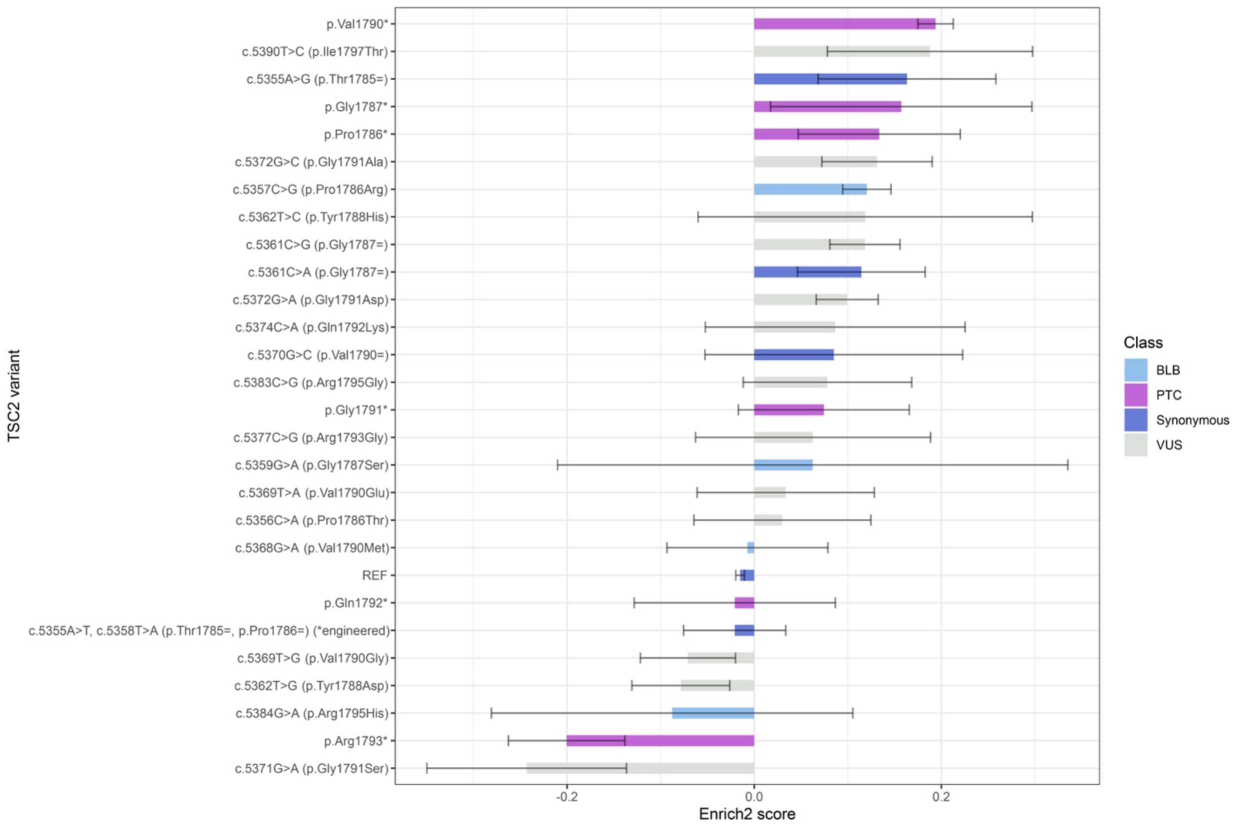

**Figure S15: Raw Enrich2 for each *TSC2* variant in exon 42.** Bars show combined enrichment score derived from 3 biological replicates. Error bars show standard error. As a reminder, exon 42 is the final exon of *TSC2*; PTC variants in this exon are not expected to cause nonsense-mediated decay of TSC2 and are expected rather to encode functional, near full-length proteins.

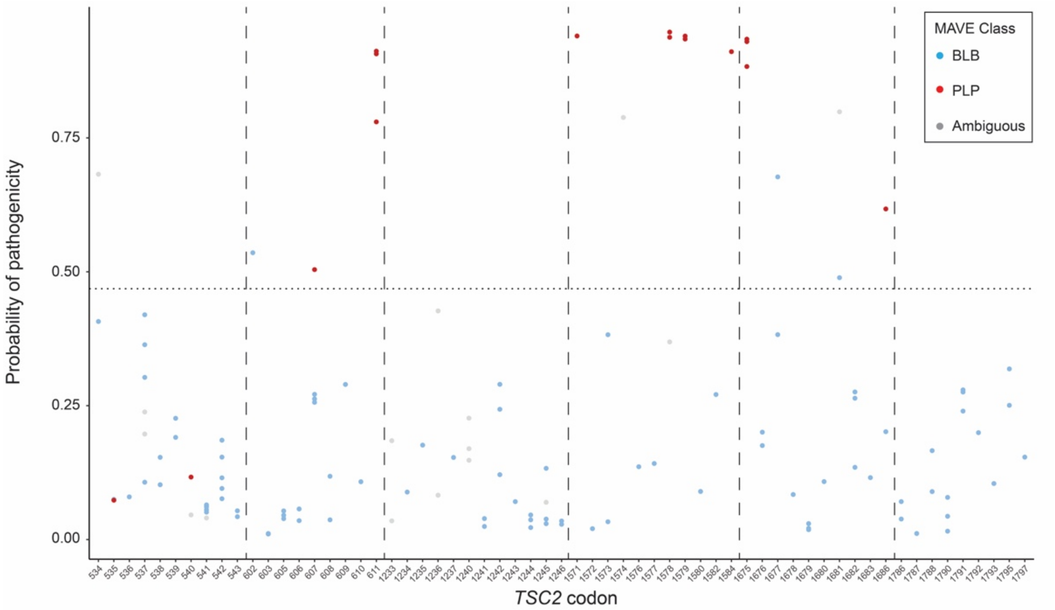

**Figure S16: TRUST predictions for each *TSC2* missense variant scored in the MAVE dataset.** Horizontal dashed line indicates the optimal threshold for binary classification of pathogenicity for TRUST. Vertical dashed lines separate each prime editing locus. Missense variants are colored by their MAVE class.

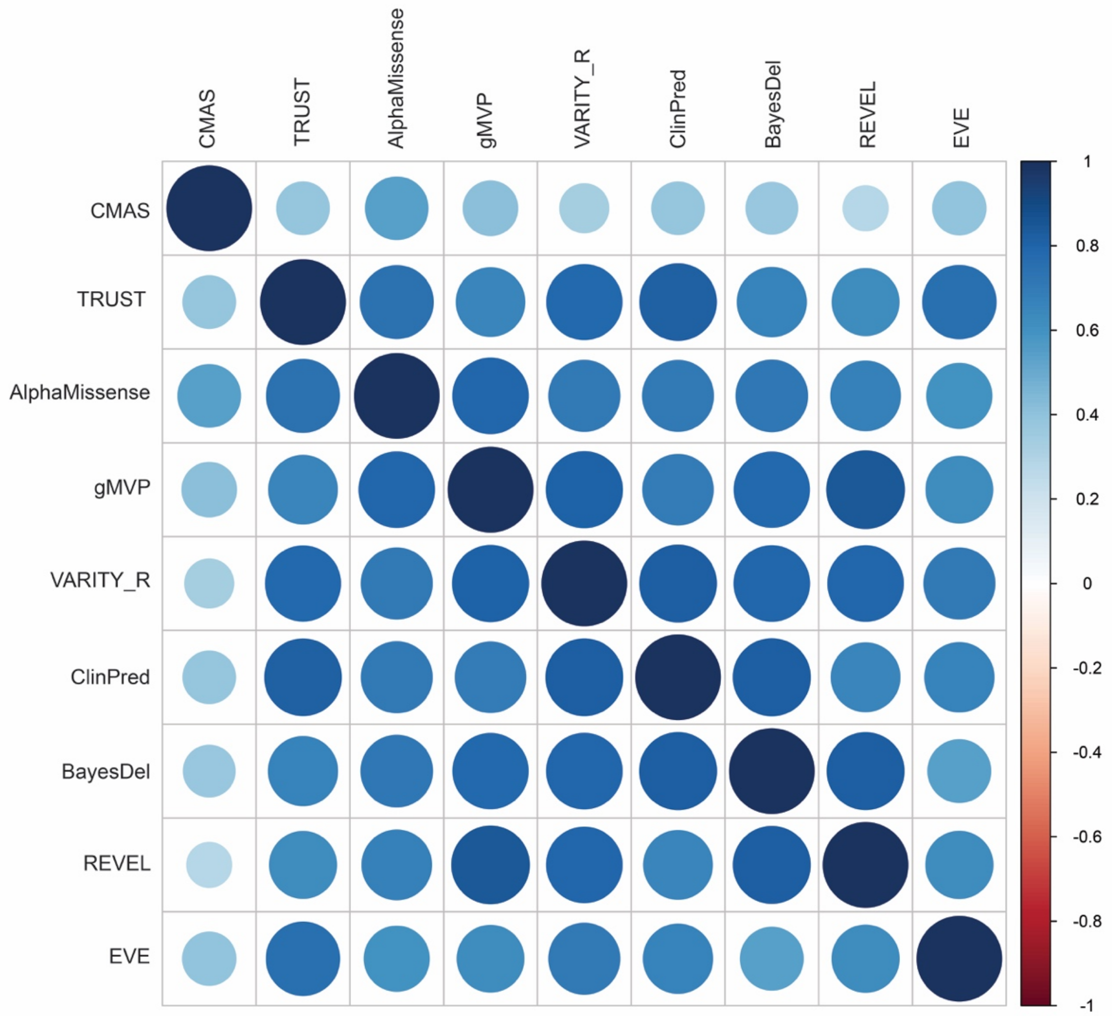

**Figure S17: Correlation plot of MAVE enrichment score, TRUST prediction, and best-performing global VEP predictions.** Scores derived using Spearman correlation metric.

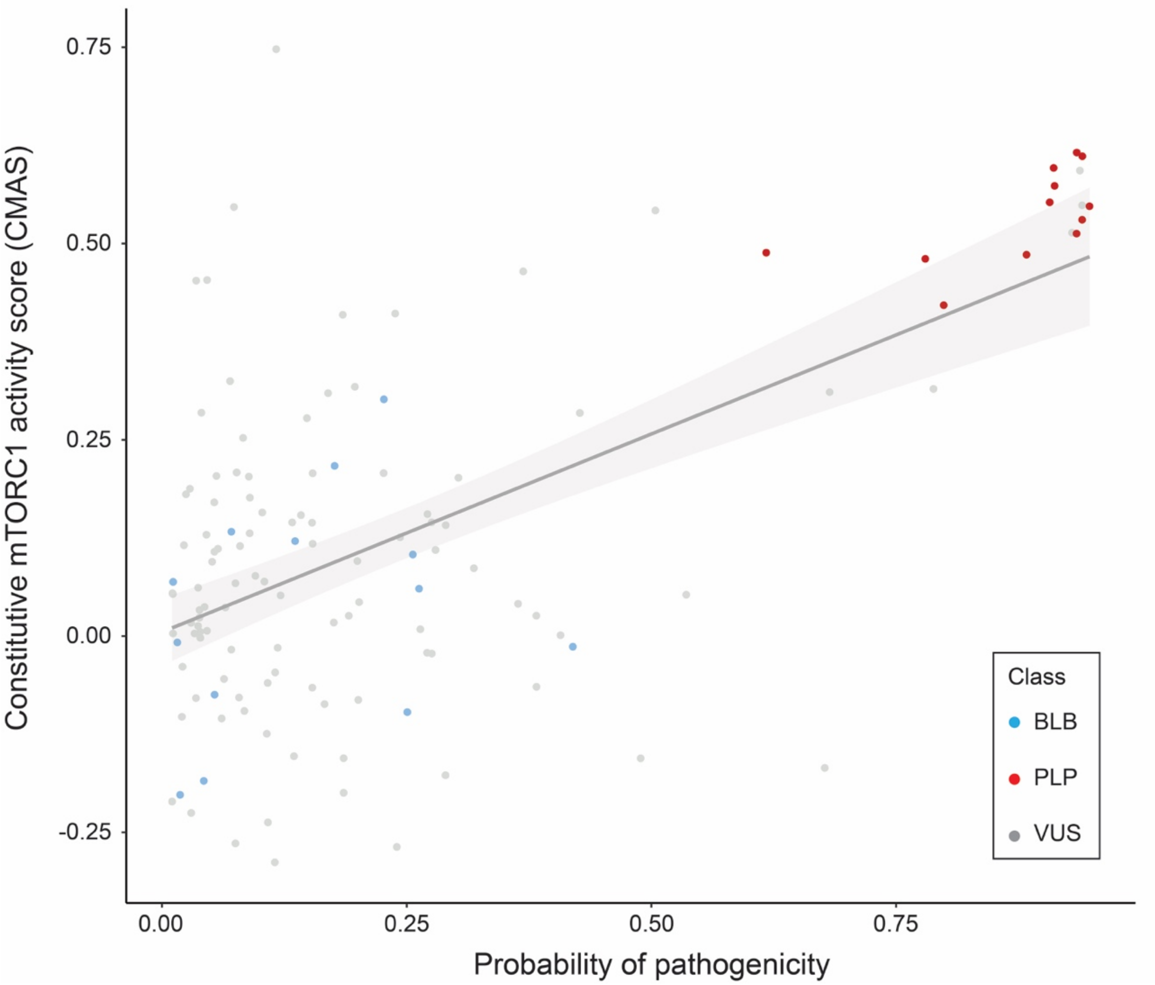

**Figure S18: Scatterplot with line of best fit for TRUST output and MAVE enrichment score.** The two are moderately correlated (r=0.604; Pearson; p < 0.001). Missense variants scored by both methods (n=114) are colored by their ClinVar class.

**Supplementary references**

dbNSFP (v4.4) was used for annotation of *TSC2* missense variants with most input features for supervised learning or *in silico* predictions from globally-trained VEPs for benchmarking.[1, 2] Other bioinformatic software packages used for *TSC2* missense annotation included: I-Mutant2.0, MAESTRO, and mCSM.[3-5]
